# Prediction of ligand-dependent conformational sampling of ABC transporters by AlphaFold3 and correlation to experimental structures and energetics

**DOI:** 10.64898/2026.02.19.706864

**Authors:** Qingyu Tang, Tianqi Wu, Brandon Soubasis, Hassane S. Mchaourab

## Abstract

AlphaFold3 architecture represented an important leap relative to Alphafold2 by enabling the inclusion of protein ligands in the prediction network. Ligand-dependent structural rearrangements are inherently difficult to predict computationally as they imply transitions between states separated by large energy differences. Here we apply AlphaFold3 to predict nucleotide-dependent changes in the conformational cycle of representative ABC transporters that have been extensively investigated by experimental structural biology techniques. We show that under similar conditions, AlphaFold3 predictions sample experimentally observed conformations. Moreover, the heterogeneity of these predictions correlates with experimental measures of dynamics obtained from multiple techniques. For couple of the tested transporters, the implied relative energetics of the conformations mirror their experimental counterpart. Remarkably, AlphaFold3 predicts previously unobserved conformations that have been implied to be sampled by ABC transporters. Finally, we report preliminary results showing that postulated sequence determinants of conformational changes modify the predictions of AlphaFold3. Although hundreds of ABC transporter structures have been determined and were included in the training data of AF3, we propose that aspects of its predictions reflect extrapolation of principles learned from these structures.

---

Protein function entails the sampling of multiple conformations along a rugged energy landscape. The enumeration of these conformations, the characterization of their relative energies, and the kinetics of their interconversion is a central task in protein science that has implications and applications in numerous fields. For decades, protein scientists have meticulously taken on this challenge often laboring under experimental conditions that obscure one or more of these conformations. The experimental bottleneck has been somewhat alleviated by the development of high-resolution single particle cryo-EM which made possible the determination of structures at the energy minima of the landscape, sometimes from the same cryo-EM sample grid(*1–5*). The emergence of deep learning protein structure prediction methods holds the promise of mapping protein conformational landscape *in silico*(*6–10*). The open codes for many of these methods along with user-friendly interfaces have sustained a surge in their applications across subfields of protein science. Here we are focused on the AlphaFold platform developed and deployed by the DeepMind team(*6*).

To generate conformational ensembles from a single input sequence using the AlphaFold2 (hereafter referred to as AF2) network(*7*), we have previously presented and benchmarked approaches to ‘hacking” AF2(*11, 12*). The original approaches exploited subsampling of the multiple sequence alignment (MSA) as well as *in silico* mutagenesis of the MSA. Subsequent robust variations on the subsampling theme have been published(*13–15*). Collectively, these methods show that AF2 can predict distinct conformations of the same protein that correspond to experimental structures. While attempts have been introduced to infer Boltzmann distributions from AF2(*16*), the implied distributions of conformations is not readily correlated with energetics.

Despite the introduction of AlphaFold3 (hereafter referred to as AF3) (*6*), the task of predictions of multiple conformations remains outstanding. As noted by its developers, AF3 does not display a robust ability to predict alternative protein conformations(*6*). However, unlike it AF2 predecessor, AF3 allows the inclusion of protein ligands in the prediction network. Conformational sampling of many proteins, including for example receptors and transporters, requires ligand binding to overcome a large energy difference between conformations. Application of MSA subsampling to predict the conformational space of such systems has not been extensively investigated although we find that for a limited set of systems, subsampling in the context of AF2 does not predict the spectrum of conformations observed under different experimental conformations. In this work, we investigate if AF3 with its diffusion architecture coupled to the introduction of ligands can overcome this limitation. We showcase the success of this approach on the ABC exporters family where conformational transitions are driven by binding and hydrolysis of ATP. ABC transporters power large structural rearrangements of their transmembrane domains (TMDs) from the energy of ATP binding and hydrolysis in the nucleotide-binding domains (NBDs) (*17–22*). More than 200 structures of ABC transporters have been determined most notably by cryo-EM revealing the conformations populated in response to the cycle of ATP binding and hydrolysis(*17, 18*).

## RESULTS

We focused our investigation on representatives of type IV ABC exporters that have been studied by a combination of structural biology techniques and represent distinct ABC exporters subfamily. MsbA, an archetype of ABC homodimers, has been extensively studied by EPR, crystallography and cryo-EM(*23–31*). The conformational cycle of the heterodimeric ABC transporter, TmrAB, was revealed in seminal work by Tempe/Moeller and colleagues(*3, 32–34*). BmrCD is another extensively investigated ABC heterodimer with distinct energy landscape as compared to TmrAB(*1, 2, 35, 36*). Both BmrCD and TmrAB have one impaired NBD where the catalytic glutamate has been replaced leading to what is referred to as a degenerate nucleotide binding site. Finally, the ABCB1 or P-glycoprotein (Pgp) transporter is a single polypeptide chain with two half transporters displaying extensive sequence divergence. Structures of Pgp have been determined from multiple organisms(*37–40*). For all the predictions reported below, AF3 had access to the template module. Predictions without templates are presented in the supplementary data and will be referenced in the appropriate context.

### MSA subsampling of ABC exporters yields limited structural variability

Because of its robustness in the context of AF2, we carried out MSA subsampling in the AF3 environment on all 4 representative ABC exporters in the absence of nucleotides. We binned the resulting 500 models for each transporter by their respective RMSDs to the corresponding experimental structures. We observed substantial conformational heterogeneity for MsbA predictions, yet it appeared that most models had large deviations from the experimental structures (Fig. 1). In contrast, while Pgp and TmrAB models were less heterogenous, they were no closer to the structures as evidenced by their RMSDs. Finally, BmrCD showed preference to sample the IF conformation with minor population of OF-like structures. We note that Pgp contains a linker regions that is presumed to be flexible hence it has not been observed experimentally(*39–41*).

**Fig. 1.**
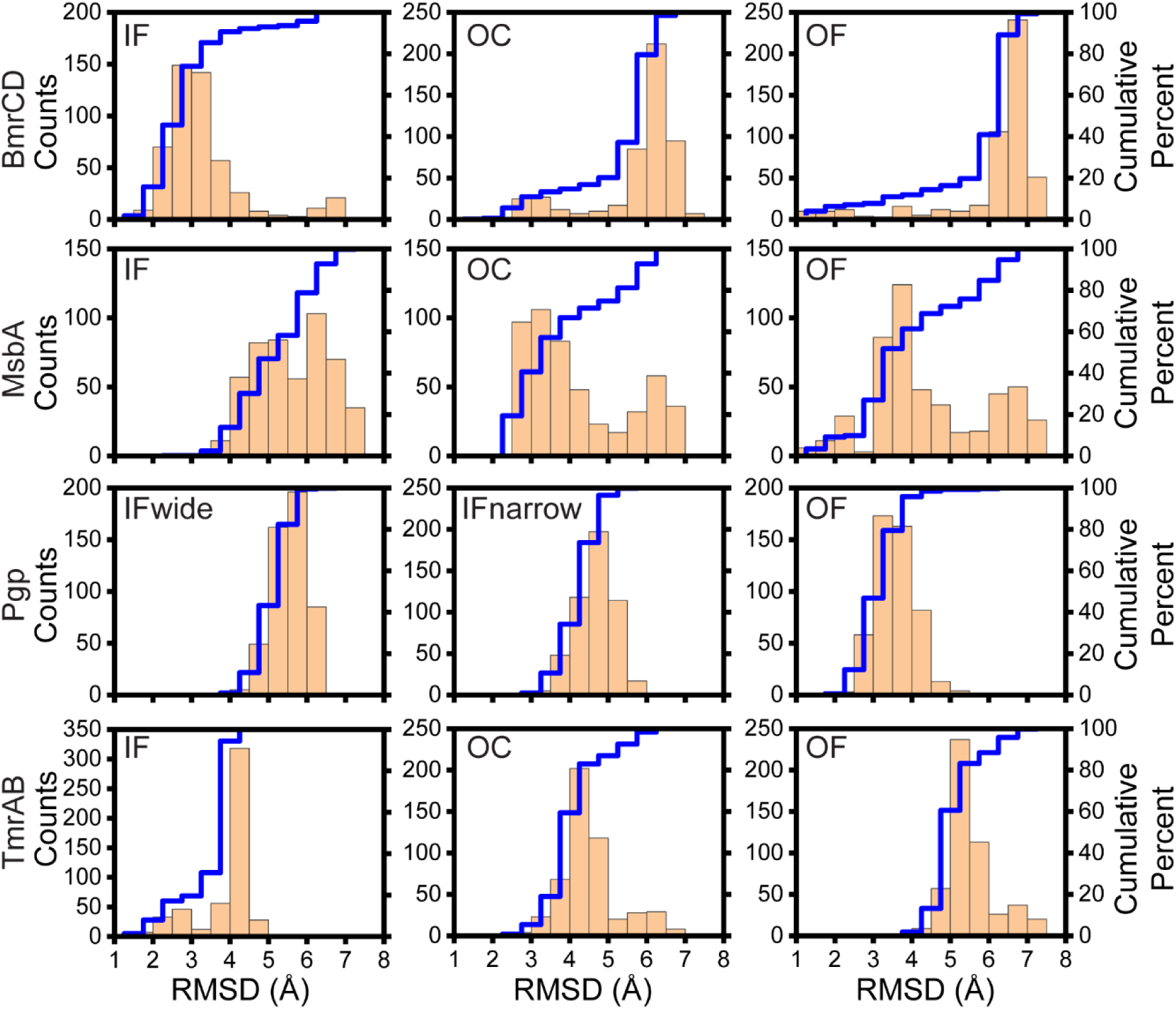
RMSD analysis of models predicted by MSA subsampling of AF3. For BmrCD, MsbA, Pgp, and TmrAB, subsampling was applied with Neff of 5 and without ligands, running for 100 iterations with random seeds. Each iteration generated 5 models. The distribution of models across conformational states for each transporter was determined by RMSD analysis using MMalign/TMalign against the reference cryo-EM structures. Reference PDB IDs: BmrCD (IF: 8FMV, OC: 8T1P, OF: 9CUS), MsbA (IF: 5TV4, OC: 5TTP, OF: 8DMM), TmrAB (IF: 6RAN, OC: 6RAI, OF: 6RAH), Pgp (IF-wide: 3G5U, IF-narrow: 6QEX, OF: 6C0V). RMSD analysis was performed with Origin software (bin size equals to 0.5 Å).

### AF3 models in the presence of nucleotides capture multiple conformations

In contrast to MSA subsampling, addition of nucleotides induced distinct and reproducible conformational preferences. The combination of nucleotides included in the predictions aimed to reproduce the experimental conditions of the available structures to the extent possible. For illustration, Figure 2 compares representative predicted AF3 models, selected by Foldseek from the ensemble of models, with the corresponding experimental structures. The figure shows that whereas AF3 successfully predicted models that are similar to the experimental structures for MsbA, and BmrCD, the OF conformation of TmrAB was conspicuously absent (Table 1). We note that Pgp’s experimental OF structure is rather occluded on the extracellular side similar to the predicted conformation.

**Fig. 2.**
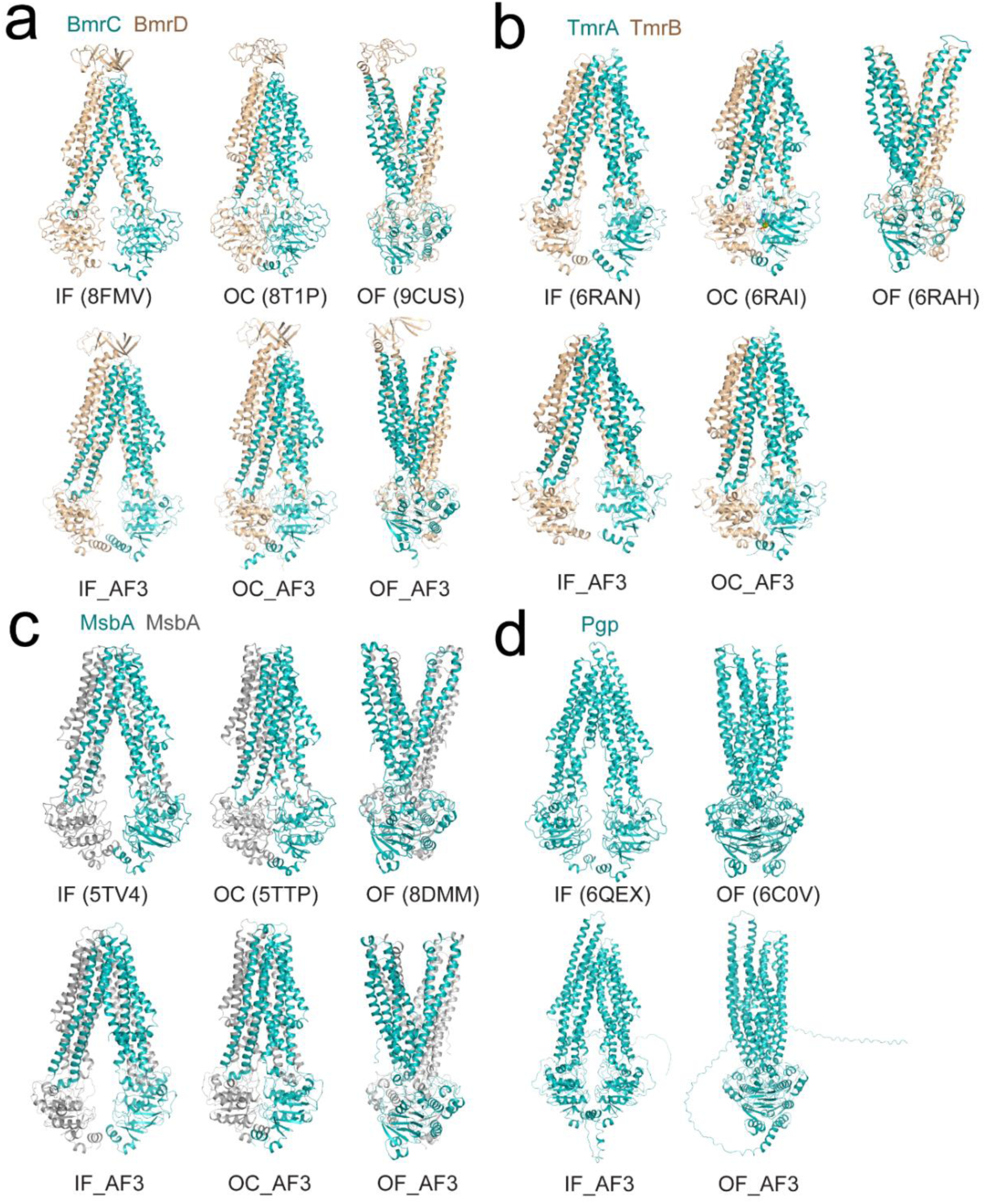
Representative models of IF, OC, and OF conformations predicted by AF3. Ribbon representations of the different conformations are shown for BmrCD (**a**), TmrAB (**b**), MsbA (**c**), and Pgp (**d**). For each transporter, the upper panels show the reference cryo-EM structures, and the lower panels show AF3 predicted models selected by Foldseek (See Methods for more details). AF3 did not predict an OF conformation for TmrAB. Relevant PDB IDs for the cryo-EM structures are indicated. Subunits BmrD, TmrB are shown in tan color, one protomer of MsbA is shown in grey color, all other subunits are shown in light sea green color. Ligands are omitted for clarity.

**Table 1.** TM-scores and RMSD values between the cryo-EM structures and representative AF3 models.

|  | **Cryo-EM structure** | **AF3 model** | **TM-score\*** | **RMSD (Å)\*** |
| --- | --- | --- | --- | --- |
| **BmrCD** | IF (8FMV) | BmrCD_IF_AF3 | 0.97 | 1.83 |
|  | OC (8T1P) | BmrCD_OC_AF3 | 0.98 | 1.67 |
|  | OF (9CUS) | BmrCD_OF_AF3 | 0.99 | 1.30 |
| **TmrAB** | IF (6RAN) | TmrAB_IF_AF3 | 0.93 | 3.17 |
|  | OC (6RAI) | TmrAB_OC_AF3 | 0.94 | 2.84 |
| **MsbA** | IF (5TV4) | MsbA_IF_AF3 | 0.93 | 3.33 |
|  | OC (5TTP) | MsbA_OC_AF3 | 0.94 | 2.76 |
|  | OF (8DMM) | MsbA_OF_AF3 | 0.99 | 1.25 |
| **Pgp** | IF (6QEX) | Pgp_IF_AF3 | 0.92 | 3.47 |
|  | OF (6C0V) | Pgp_OF_AF3 | 0.98 | 1.62 |

Note: \* TM-score and RMSD were calculated by MM-align(51) (for BmrCD, TmrAB, and MsbA) and TM-align(52) (for Pgp).

More quantitatively, Figures 3 and 4 report a RMSD probability histogram that emphasizes how inclusion of nucleotides shift the predictions of AF3 (Figs. S1 and S2). Analysis of these changes, however, further illustrates stark differences between the predictions for the four ABC exporters under different nucleotide and magnesium combinations. For BmrCD, we observe that the models for Apo and 2Mg^2+^/2ATP yield high probability and low RMSD for the IF and OF conformations respectively, as expected. In contrast, we were unable to find nucleotide combinations that force AF3 to predict the OF conformation of TmrAB (Fig. S1). The results for BmrCD and TmrAB are notable considering that the latter had an OF structure in the PDB(*3*), i.e. available for the template module, whereas the former did not.

**Fig. 3.**
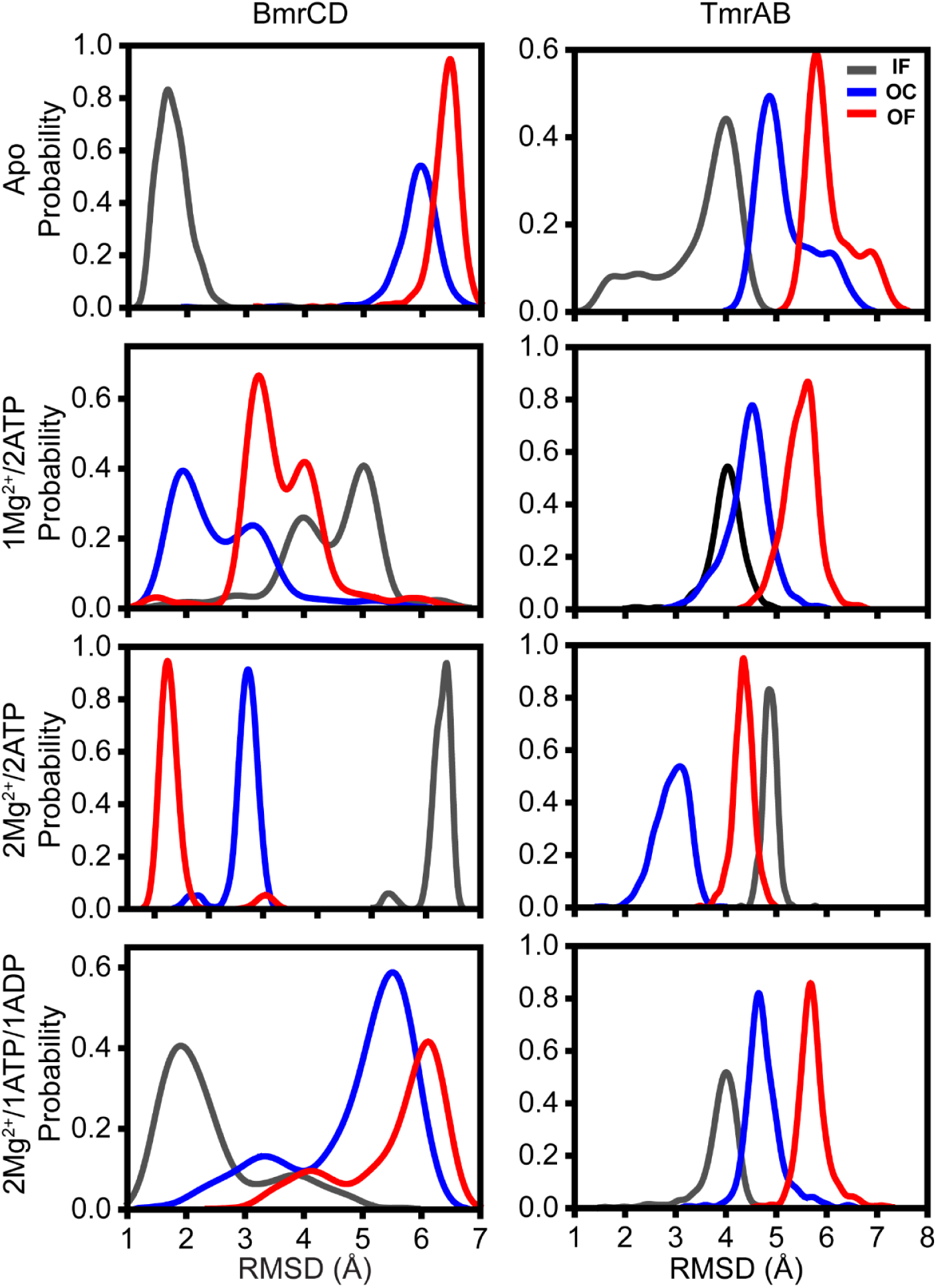
RMSD analysis of AF3 predictions for BmrCD and TmrAB in the presence of combinations of nucleotide and Mg^2+^. AF3 predictions were performed for BmrCD and TmrAB under four nucleotide conditions: apo, 1Mg^2+^/2ATP, 2Mg^2+^/2ATP, and 2Mg^2+^/1ATP1ADP. For each condition, 500 models were generated using random seeds. RMSD values for the models in each condition were calculated with MMalign by aligning them to the reference cryo-EM structures of IF, OC, and OF conformations. The resulting RMSD distributions for the IF, OC, and OF are shown in grey, blue and red, respectively. Reference PDB IDs: BmrCD (IF: 8FMV, OC: 8T1P, OF: 9CUS), TmrAB (IF: 6RAN, OC: 6RAI, OF: 6RAH).

**Fig. 4.**
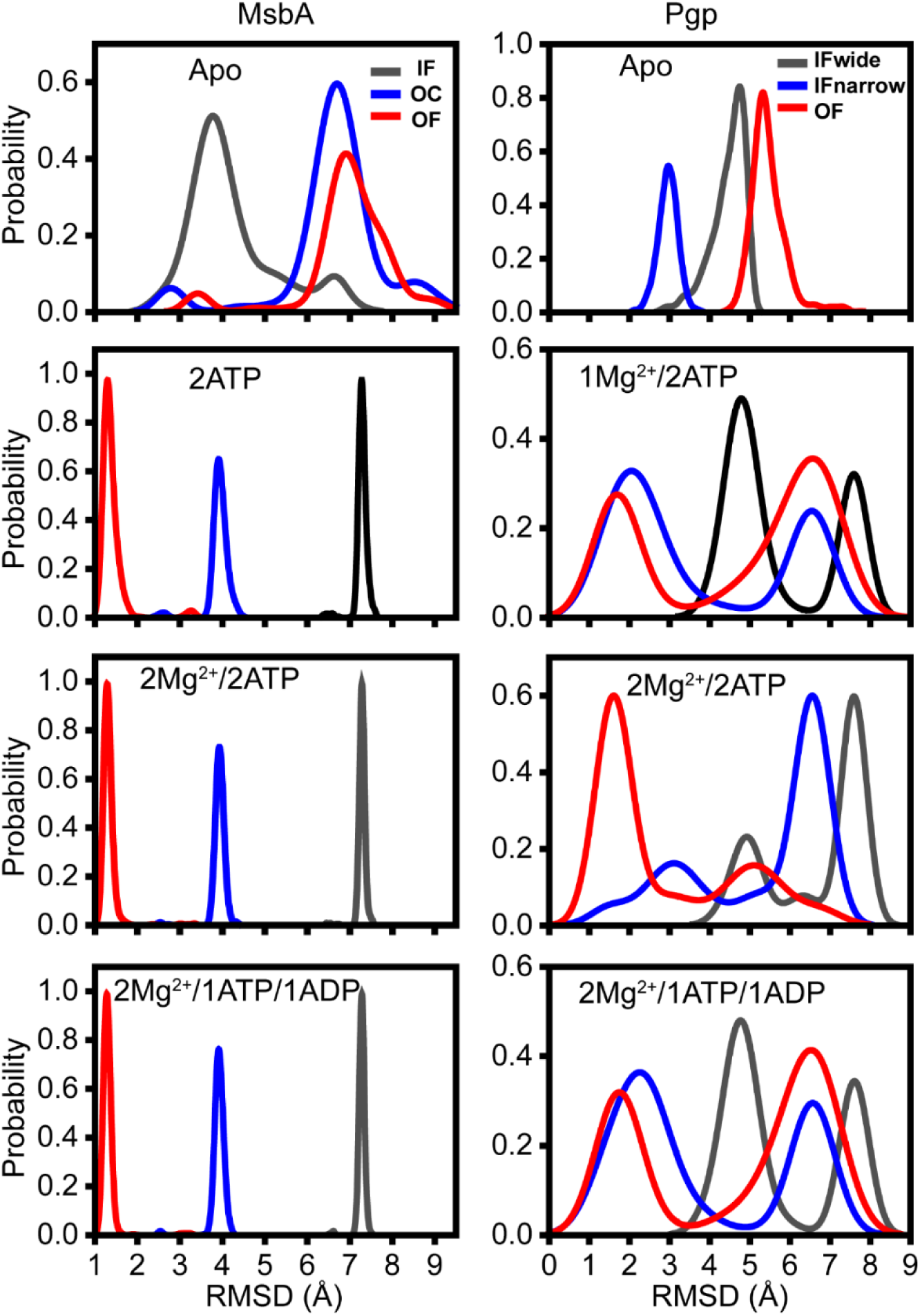
RMSD analysis of AF3 predictions for MsbA and Pgp in the presence of combinations of nucleotide and Mg^2+^. AF3 predictions were performed for MsbA under apo, 2ATP, 2Mg^2+^/2ATP, and 2Mg^2+^/1ATP1ADP conditions, and for Pgp under apo, 1Mg^2+^/2ATP, 2Mg^2+^/2ATP, and 2Mg^2+^/1ATP1ADP conditions. For each condition, 500 models were generated using random seeds. RMSD values for the models in each condition were calculated by aligning them to the reference cryo-EM structures of IF, OC, and OF conformations using MMalign for MsbA and TMalign for Pgp. The resulting RMSD distributions for the IF, OC, and OF are shown in grey, blue and red, respectively. Reference PDB IDs: MsbA (IF: 5TV4, OC: 5TTP, OF: 8DMM), Pgp (IF-wide: 3G5U, IF-narrow: 6QEX, OF: 6C0V).

Models for the Apo condition of MsbA yielded broad distributions that primarily reflect the IF conformation. Remarkably inclusion of two nucleotide molecules, including 2Mg^2+^/2ATP or 2Mg^2+^/1ATP1ADP, in different combinations consistently shifted the conformational preference to the OF conformation (Fig. S2). This shift was accompanied by a tighter RMSD distributions suggesting a highly homogenous ensemble of models. In contrast, Pgp models displayed heterogenous RMSDs with the nucleotide-free and 2Mg^2+^/2ATP conditions favoring the IF and OF conformations respectively (Fig. S2).

Other combinations of nucleotides increased the heterogeneity of the predictions (Figs. S1 and S2). We simulated conditions under which only one Mg^2+^ was included along with two molecules of ATP. Except for MsbA which appeared to favor OF conformation under multiple nucleotide conditions, the other transporters showed variable conformational preferences. For BmrCD, the predictions partly favored the OC conformation whereas for Pgp a population of OF conformation (which is essentially an OC) was detected along with a population of IF conformation. Notably, AF3 predicted an OF conformation for MsbA in the presence of 2Mg^2+^/2ADP which is inconsistent with experimental data(*28*) (Fig. S3). Excluding templates did not change the prediction under these conditions (Fig. S3). Overall, for all transporters except the homodimer MsbA, a single Mg^2+^ triggered changes in the preferred conformation suggesting a pivotal role in the cycle.

### Predicted ensemble heterogeneity is consistent with measured DEER implied heterogeneity

To compare the predicted models with the conformational ensembles deduced from previously published DEER data, we measured the Cα distances at the extracellular and intracellular sides of the transporters. For BmrCD, MsbA and Pgp, the residues were selected based on the experimental spin label pairs(*1, 23, 35, 36, 42, 43*). These pairs have been shown to report the IF to OF transition in the presence of various nucleotides and substrates (Fig 5, solid traces). Figure 6 and Figs. S4-S5 show the Cα-Cα / Cβ−Cβ distance distributions at the extracellular and intracellular pairs obtained from 500 predicted models under different nucleotide and Mg^2+^ conditions. Addition of 2Mg^2+^/2ATP induces opposite distance changes at the two sides of these transporters reflecting transition between IF and OF conformations. However, the distributions reveal that the apparent energetic preferences, as captured by the ratio of these populations in the presence of 2Mg^2+^/2ATP, differ between transporters. MsbA and BmrCD are almost completely switched to the OF conformation. On the other hand, Pgp continues to sample the IF conformation. Notably for MsbA, the first ATP binding induces a lager distance distribution at the extracellular side (158/158) and a shorter distance distribution at the intracellular side (103/103). The conformational shift is completed upon the second ATP binding (Fig. 6),

**Fig. 5.**
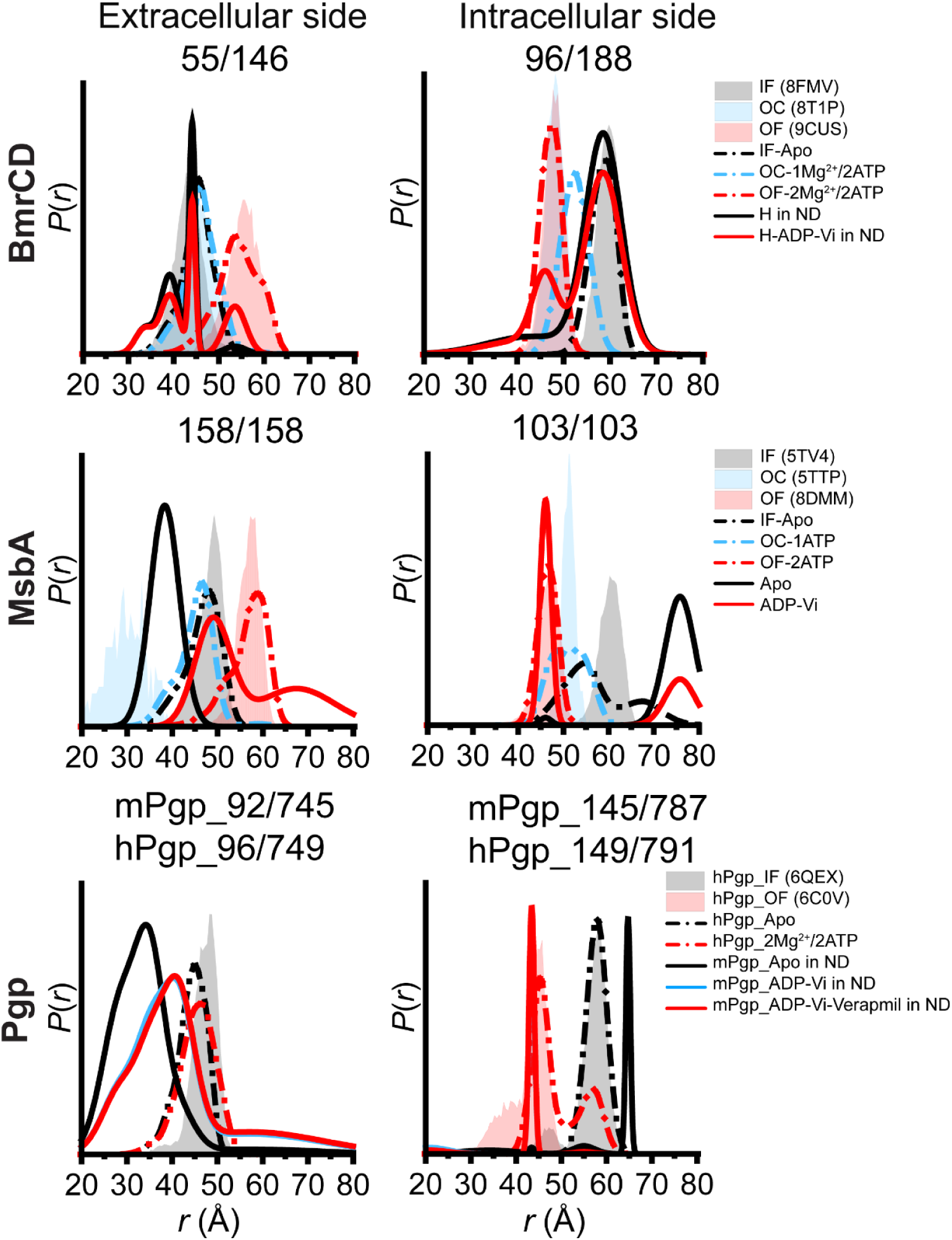
Model validation using cryo-EM structures and DEER spectroscopy. Predicted models were validated by comparing simulated distance distributions to experimental DEER data for the spin-labeling sites on the extracellular (left panels) and intracellular (right panels) sides of BmrCD, MsbA, and Pgp. Spin-labeled pairs 55/146 and 96/188 are denoted for 55^BmrC^/146^BmrD^ and 96^BmrC^/188^BmrD^, respectively. For each pair, the shaded area represents the distribution simulated from cryo-EM structure using MDDS. Residue 96 was not visible in cryo-EM structure of hPgp (PDB ID: 6C0V), precluding prediction of distance distributions. The dashed line shows the distribution simulated from the corresponding ChiLife models, and the solid line shows the experimental DEER distribution.

**Fig. 6.**
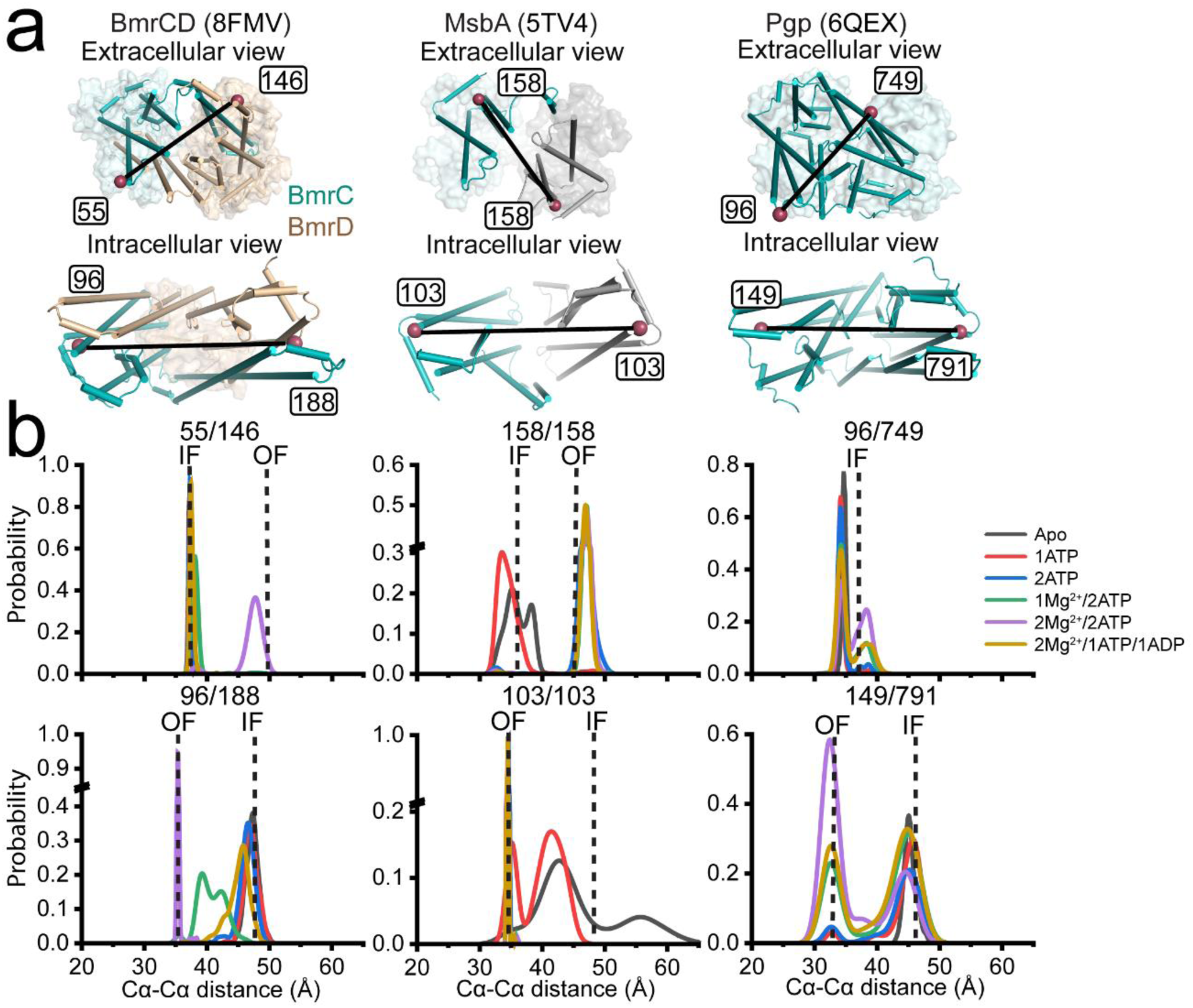
Distinct conformational transitions revealed by the Cα-Cα distance analysis in BmrCD, MsbA, and Pgp. **(a)** Close-up views highlight the residue pairs on the extracellular side (upper panels) and intracellular side (lower panels) of BmrCD (left panels; PDB ID: 8FMV), MsbA (middle panels; PDB ID: 5TV4), and Pgp (right panels; PDB ID: 6QEX). Transmembrane domain (TMD) helices are shown as cylinders; NBDs or ECD (in BmrCD) are shown in transparent surfaces. The residue pairs are indicated with raspberry spheres. The color scheme is consistent with Fig. 2. **(b)** Distribution plots of the Cα-Cα distances of the residue pairs on the extracellular side (upper panels) and intracellular side (lower panels) of BmrCD (left panels), MsbA (middle panels), and Pgp (right panels). Data for six ligand conditions (apo, 1ATP, 2ATP, 1Mg^2+^/2ATP, 2Mg^2+^/2ATP, and 2Mg^2+^/1ATP1ADP) are plotted for each pair. The dashed lines indicate the distance derived from the cryo-EM structures of the corresponding protein in its IF and OF conformations. Distribution analysis was performed with Origin software (bin size equals to 1 Å; data height is calculated by Relative Frequency).

Other nucleotide conditions shown in Figure 6 further expose apparent energetic differences. On one extreme, MsbA is in a OF conformation under all conditions except apo and in the presence of 1 ATP where MsbA partially samples an occluded conformation (Fig. S6). In contrast except for the 1Mg^2+^/2ATP, BmrCD is in an occluded conformation suggesting that binding of the second Mg^2+^ induces the transition to the OF conformation as experimentally observed(*1*). Finally, Pgp appears to partition primarily between the IF and OF conformations even in the presence of 2Mg^2+^/2ATP. This is consistent with the observation that the OF conformation is rarely sampled.

### AF3 predicts previously unobserved conformations of BmrCD

RMSD and distance analyses predict conformations that have not been observed experimentally. This is illustrated by the green trace for the BmrCD 96/188 pair in Figure 6 in the presence 1Mg^2+^/2ATP. RMSD of the 500 models relative to the experimental OC and OF conformations show that the major population of predicted models are close to the former (Fig. 3). However, the second major population, which has a RMSD of 3 Å compared to OC structure (PDB 8T1P) (Figs. 7 and S6, Tables 2 and 3), hereafter referred to as asymmetric OC, deviates on the intracellular side yielding the green distance population in Figure 6. Remarkably, inspection of the nucleotide binding sites (NBSs) in the asymmetric OC conformation reveals preference for Mg^2+^ binding at the degenerate site (Fig. S7a) which is disengaged in a fraction of the models (Stage I) whereas the consensus site is engaged across the population of models (Fig. S7b). The distance between the ABC motif and ATP at the degenerate site increased from 3 Å to 10 Å, compared to almost no changes on average at the consensus site (Figs. 7b and S7). This conclusion is illustrated by the distance distribution observed at the NBSs. Here we monitored pairs of residues that previously reported asymmetric conformations as observed by DEEER of spin labeled pairs (*1, 35*) (Figs. S8 and S9). Thus, the transition from OC to asymmetric OC results in disengagement at the degenerate NBS site.

**Fig. 7.**
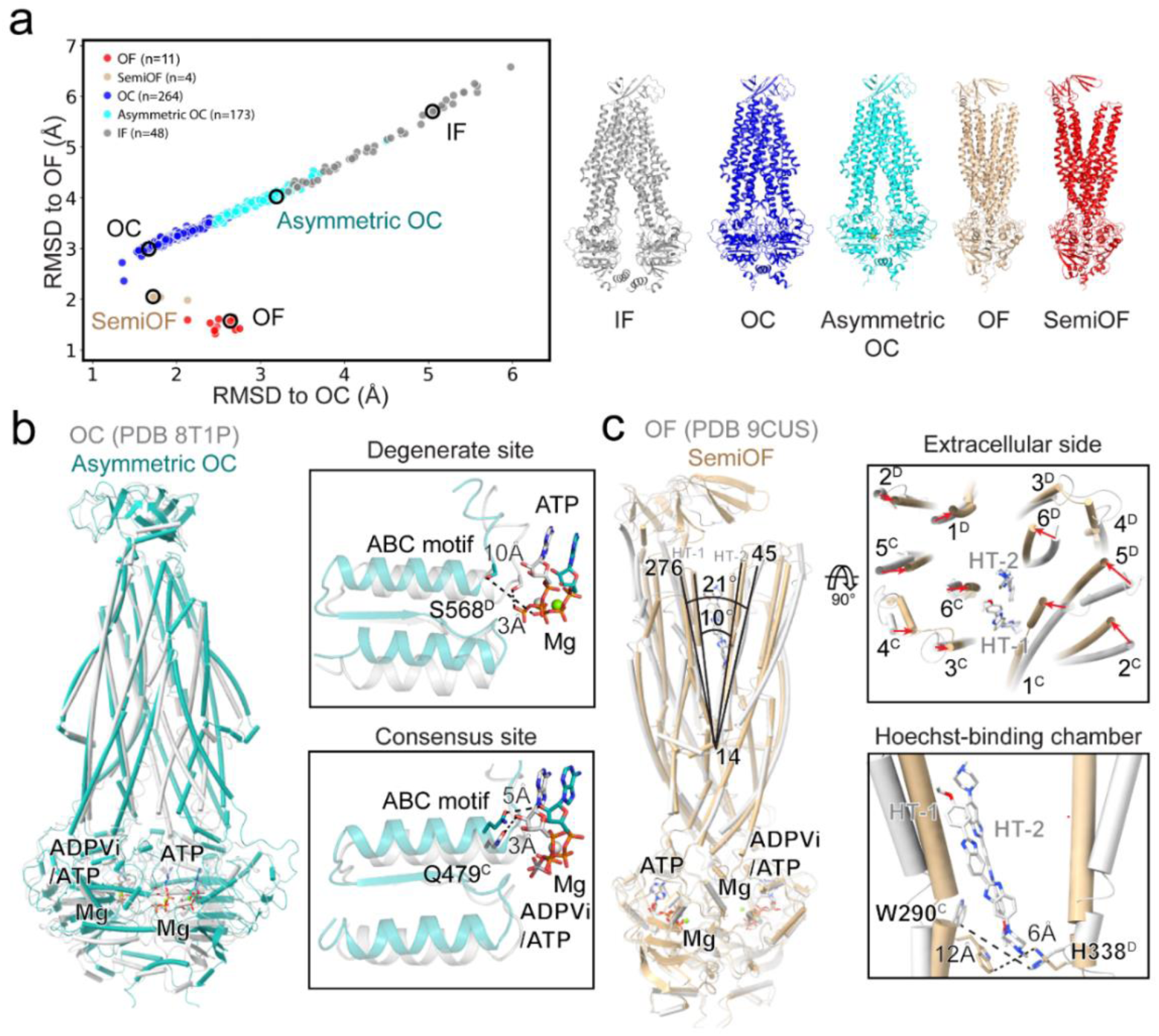
Intermediate states of BmrCD sampled out by AF3. (**a**) RMSD plot for the 1Mg^2+^/2ATP condition of BmrCD. Conformations were classified into five groups based on Cα-Cα distances analysis of the extracellular and intracellular sides: IF (grey), OC (blue), Asymmetric OC (cyan), OF (red), and SemiOF (tan). The representative model for each group selected by Foldseek is highlighted with a black circle and shown as cartoons in the right panels with Mg^2+^ depicted as spheres and ATP as sticks. **(b)** Comparison of the asymmetric OC and OC. Left panel: Superposition of the two structures. Right panels: Distances between the ABC motifs and ATP at the degenerate NBS (upper), and distances between the ABC motifs and ATP or ADPVi at the consensus NBS (lower) in the asymmetric OC and OC conformations. Relevant distances are indicated, and ligands are labeled with Mg^2+^ depicted as spheres and ATP as sticks. Asymmetric OC and OC conformations are shown in light sea green and grey, respectively. **(c)** Comparison of the semiOF and OF conformations. Left panel: Superposition of the semiOF and OF conformations aligned on BmrD reveals a smaller opening angel at the extracellular side in the semiOF. Ligands and the residues used for measurement are labeled. Upper-right panel: A top view of the extracellular side of TMD shows tighter closure of the helices in the semiOF, with large movements indicated by red arrows. Lower-right panel: Close-up view of the Trp290^BmrD^-His338 ^BmrD^ (WH) latch. The distance of W290 and H338 in the semiOF (6 Å) is similar to that in the two Hoechsts-bound, inward-facing conformation (7 Å, PDB ID: 8FMV). Relevant distances are indicated and Mg^2+^ is depicted as spheres and ATP as sticks. SemiOF and OF conformations are shown in tan and grey, respectively.

**Table 2.**
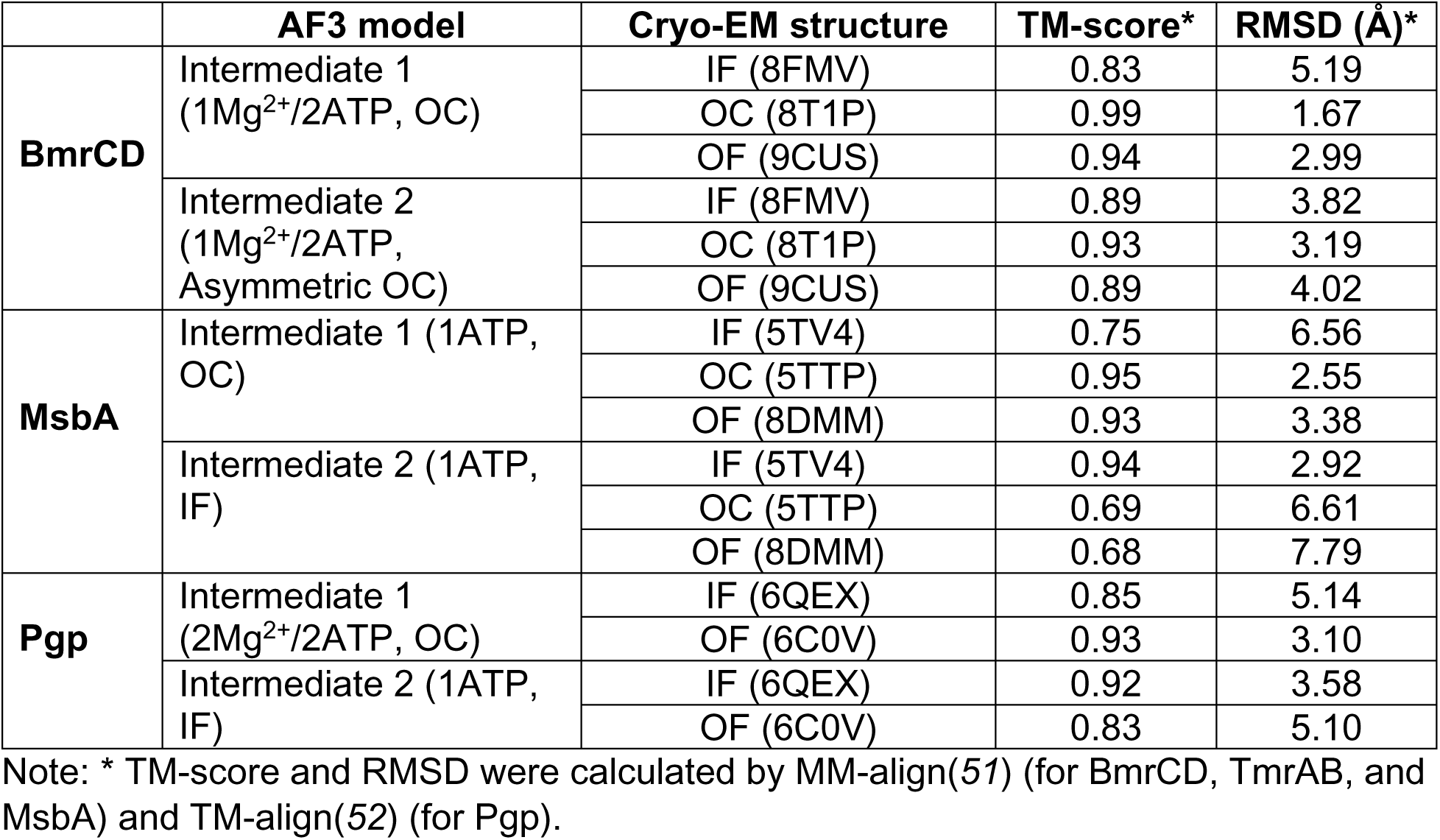
TM-scores and RMSD values between the cryo-EM structures and intermediate AF3 models.

|  | **AF3 model** | **Cryo-EM structure** | **TM-score\*** | **RMSD (Å)\*** |
| --- | --- | --- | --- | --- |
| **BmrCD** | Intermediate 1<br>(1Mg<sup>2+</sup>/2ATP, OC) | IF (8FMV) | 0.83 | 5.19 |
|  |  | OC (8T1P) | 0.99 | 1.67 |
|  |  | OF (9CUS) | 0.94 | 2.99 |
|  | Intermediate 2<br>(1Mg<sup>2+</sup>/2ATP, Asymmetric OC) | IF (8FMV) | 0.89 | 3.82 |
|  |  | OC (8T1P) | 0.93 | 3.19 |
|  |  | OF (9CUS) | 0.89 | 4.02 |
| **MsbA** | Intermediate 1 (1ATP, OC) | IF (5TV4) | 0.75 | 6.56 |
|  |  | OC (5TTP) | 0.95 | 2.55 |
|  |  | OF (8DMM) | 0.93 | 3.38 |
|  | Intermediate 2 (1ATP, IF) | IF (5TV4) | 0.94 | 2.92 |
|  |  | OC (5TTP) | 0.69 | 6.61 |
|  |  | OF (8DMM) | 0.68 | 7.79 |
| **Pgp** | Intermediate 1<br>(2Mg<sup>2+</sup>/2ATP, OC) | IF (6QEX) | 0.85 | 5.14 |
|  |  | OF (6C0V) | 0.93 | 3.10 |
|  | Intermediate 2 (1ATP, IF) | IF (6QEX) | 0.92 | 3.58 |
|  |  | OF (6C0V) | 0.83 | 5.10 |

Note: \* TM-score and RMSD were calculated by MM-align(51) (for BmrCD, TmrAB, and MsbA) and TM-align(52) (for Pgp).

**Table 3.**
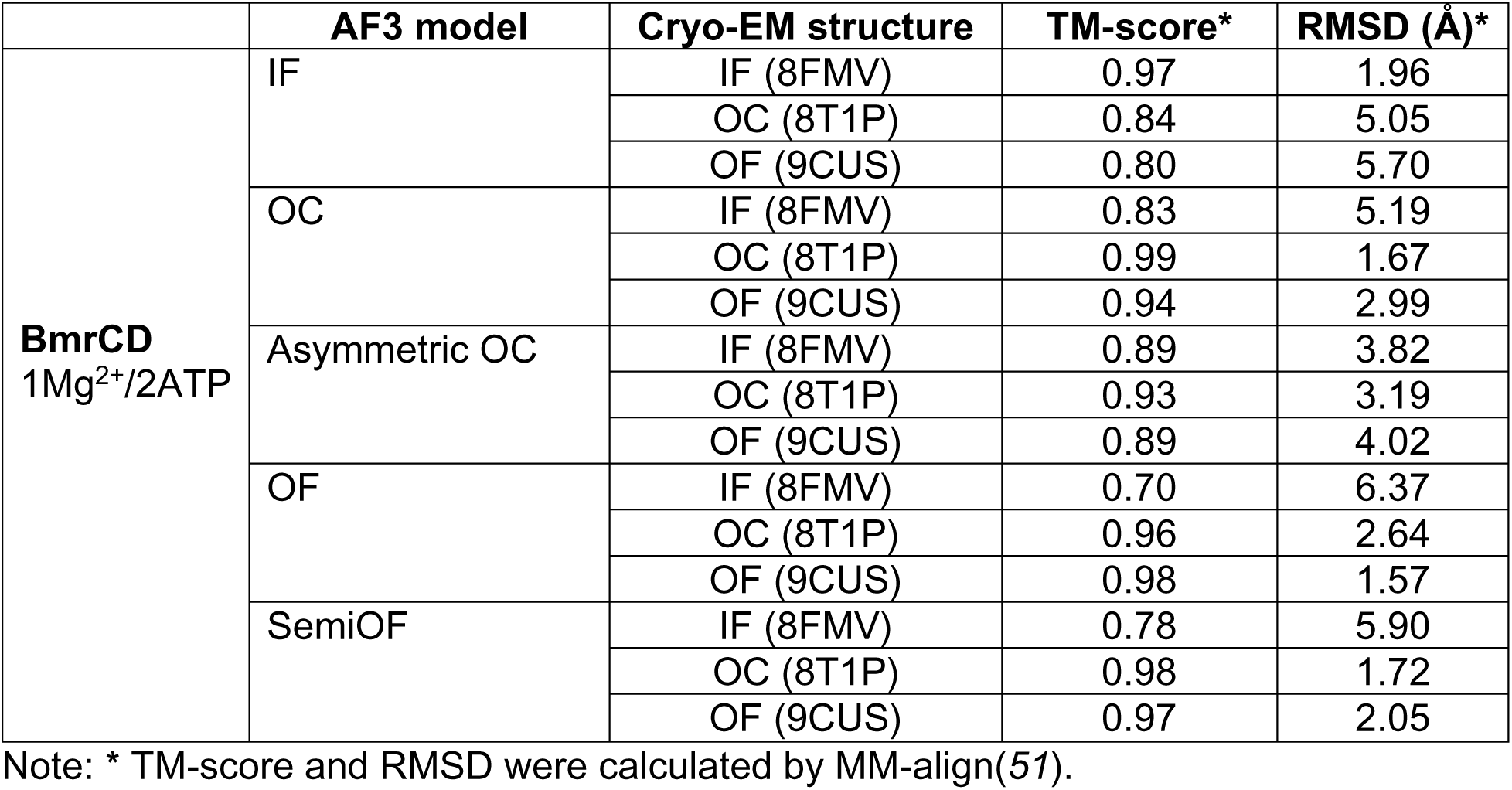
TM-scores and RMSD values between the cryo-EM structures and intermediate AF3 models of BmrCD.

|  | AF3 model | Cryo-EM structure | TM-score* | RMSD (Å)* |
| --- | --- | --- | --- | --- |
| **BmrCD**<br>1Mg<sup>2+</sup>/2ATP | IF | IF (8FMV) | 0.97 | 1.96 |
|  |  | OC (8T1P) | 0.84 | 5.05 |
|  |  | OF (9CUS) | 0.80 | 5.70 |
|  | OC | IF (8FMV) | 0.83 | 5.19 |
|  |  | OC (8T1P) | 0.99 | 1.67 |
|  |  | OF (9CUS) | 0.94 | 2.99 |
|  | Asymmetric OC | IF (8FMV) | 0.89 | 3.82 |
|  |  | OC (8T1P) | 0.93 | 3.19 |
|  |  | OF (9CUS) | 0.89 | 4.02 |
|  | OF | IF (8FMV) | 0.70 | 6.37 |
|  |  | OC (8T1P) | 0.96 | 2.64 |
|  |  | OF (9CUS) | 0.98 | 1.57 |
|  | SemiOF | IF (8FMV) | 0.78 | 5.90 |
|  |  | OC (8T1P) | 0.98 | 1.72 |
|  |  | OF (9CUS) | 0.97 | 2.05 |

Note: \* TM-score and RMSD were calculated by MM-align(51).

Furthermore, we identified 4 models in the ensemble that are OF-like (hereafter referred to as semi-OF) with RMSD values to the OF (PDB 9CUS) of around 2 Å (Fig. 7c). Superposition of these semi-OF model on the experimental OF structures (aligning the BmrD subunit) reveals that all helices on the extracellular side shifted toward the middle by approximately 2 to 7 Å, while the intracellular side showed virtually no changes (Fig. 7c). As a result, the angle between the extracellular sides of TM1 and TM6 of BmrC decreased from 21° to 10° in the semi-OF models (Fig. 7c), thereby reducing the chamber volume from 4995 Å^3^ to 2239 Å^3^ (*44, 45*). If Hoechst molecules are docked into their position in the experimental OF structure, severe steric hindrance with TM1 and TM6 of BmrC, as well as TM6 of BmrD, would result in the semi-OF confirmation suggesting that this model may be on the reset path from the OF to the IF following drug release.

### Shifts in AF3 conformational preferences is coupled to changes in heterogeneity

We noted above (Figs. 3 and 4, S1 and S2) that the RMSD distributions between predicted models and structures displayed variable widths under different conditions. To further dissect this property and relate it to the cryo-EM experimental data, we calculated the residue mean square root fluctuations (RMSF) to quantify flexibility across the predicted ensembles. All models were aligned to a reference structure (the first PDB file under each condition) using a least-squares fitting algorithm (as implemented in the Superimposer module from BioPython(*46*)) to minimize the Cα atom positional deviations. Subsequently, the RMSF was calculated based on this structural alignment. The experimental B-factors, which were determined during the refinement of the cryo-EM reference structure, were mapped onto the molecular structure and visualized using the B-factor putty representation in PyMOL (Fig. 8, red regions indicate higher flexibility).

**Fig. 8.**
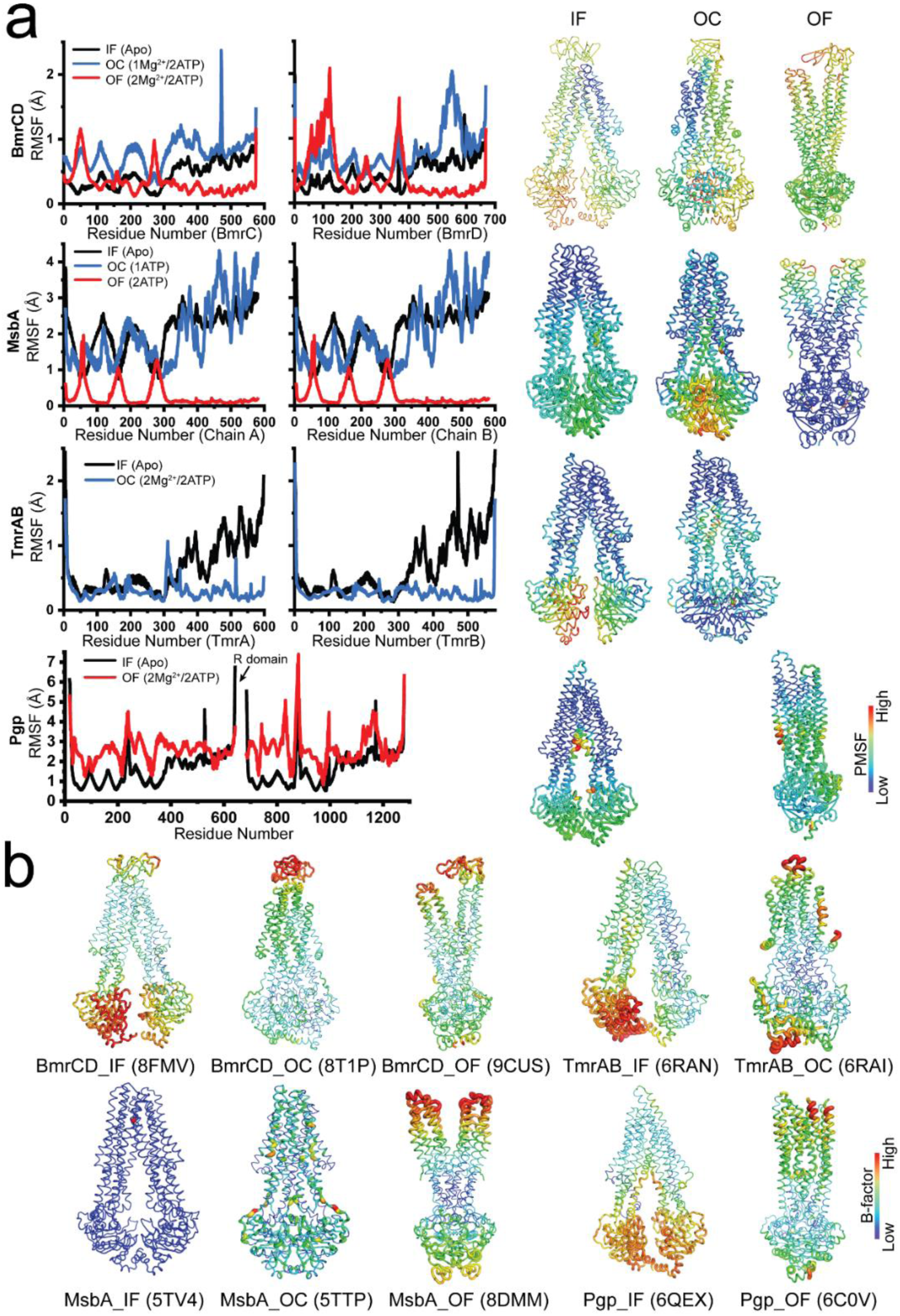
Structural fluctuation evaluated by RMSF analysis of predicted models. **(a)** RMSF plots (left panels) and their putty representations (right panels) for different conformations of BmrCD, MsbA, TmrAB, and Pgp. The conditions with dominant IF, OC, or OF conformations are selected for analysis. Both BmrCD and MsbA sampled IF, OC, and OF conformations. TmrAB sampled only IF and OC while Pgp sampled only IF and OF confirmations. **(b)** Experimental B-factor putty representations from cryo-EM structures. The B-factors from the corresponding experimental cryo-EM structures are shown for comparison. The structures are the same as those in Fig. 2.

We observed a consistent pattern between predicted and experimental fluctuations. Fluctuations in the NBDs across IF models qualitatively mirror the high B-factors in the corresponding experimental IF conformations, for BmrCD, TmrAB, and Pgp. Conversely, the extracellular regions of the TMDs exhibit greater fluctuations in the OF models, which correlates with higher B-factors in the experimental OF conformations, as observed for BmrCD and MsbA (Fig. 8).

### Coupling helices in BmrCD and TmrAB are determinants of conformational selection

The failure of AF3 to predict the OF conformation of TmrAB despite its availability in the PDB (Fig. S5), and thus in the template module of AF3, stands in stark contrast to the success in modeling BmrCD’s OF conformation which had not been deposited in the PDB when these predictions were obtained. To further investigate the origin of this finding, we targeted regions of the transporters that have been implicated in the coupling of conformational changes between the NBD and TMD, specifically the coupling helices(*47, 48*).

For this purpose, we predicted models in the presence of 2Mg^2+^/2ATP for chimeric proteins of BmrCD and TmrAB wherein the coupling helices were interchanged. The impaired catalytic sites (also referred to as degenerate sites) of TmrAB and BmrCD, i.e. where the catalytic Glutamte is replaced, are located in TmrB and BmrC respectively. Thus, the coupling helices were swapped between TmrB and BmrC, with a similar exchange between TmrA and BmrD. Specifically, 12 residues were selected for CH1 (between TM2 and TM3), and 11 for CH2 (between TM4 and TM5) (Table 4).

**Table 4.** Coupling helices switch of BmrCD and TmrAB.

|  | TmrA_CH1 (A1)/<br>BmrC_CH1 (C1) | TmrA_CH2 (A2)/<br>BmrC_CH2 (C2) | TmrB_CH1 (B1)/<br>BmrD_CH1 (D1) | TmrB_CH2 (B2)/<br>BmrD_CH2 (D2) | Rest part |
| --- | --- | --- | --- | --- | --- |
| **WT** | TMSPPFYEKNRT | SGVRVIRAYVQ | KMPIRYFDNLPA | QGMTIIQAFRH | BmrCD |
| **CD12BA** | TLDRDFYHKHRV | SGIRVVKGYAL | RLHPGFYDRNPV | SGVETIQLFVK | BmrCD |
| **CD12BA-C1** | TMSPPFYEKNRT | SGIRVVKGYAL | RLHPGFYDRNPV | SGVETIQLFVK | BmrCD |
| **CD12BA-C2** | TLDRDFYHKHRV | SGVRVIRAYVQ | RLHPGFYDRNPV | SGVETIQLFVK | BmrCD |
| **CD12BA-D1** | TLDRDFYHKHRV | SGIRVVKGYAL | KMPIRYFDNLPA | SGVETIQLFVK | BmrCD |
| **CD12BA-D2** | TLDRDFYHKHRV | SGIRVVKGYAL | RLHPGFYDRNPV | QGMTIIQAFRH | BmrCD |

Compared to predictions of wild-type (WT) BmrCD which yields 95% OF conformation, the engineered BmrCD with all four coupling helices replaced by those of TmrAB (referred as BmrCD12BA) displayed 100% OC conformation (Figs. 9 and S10). To pinpoint the crucial coupling helices, predictions with four CH replacements were performed, which restored varying populations of OF conformation (Fig. 9). We observed that the replacement of a CH with evolutionary distant CH yielded a larger population of OC conformation (Fig. 9d). Finally, the chimera wherein wherein the CH pf BmrC were replaced by TmrA’s and BmrD’s were replaced by TmrB’s (BmrCD12AB) yielded less pronounced effects compared to BmrCD12BA (Fig. 9).

**Fig. 9.**
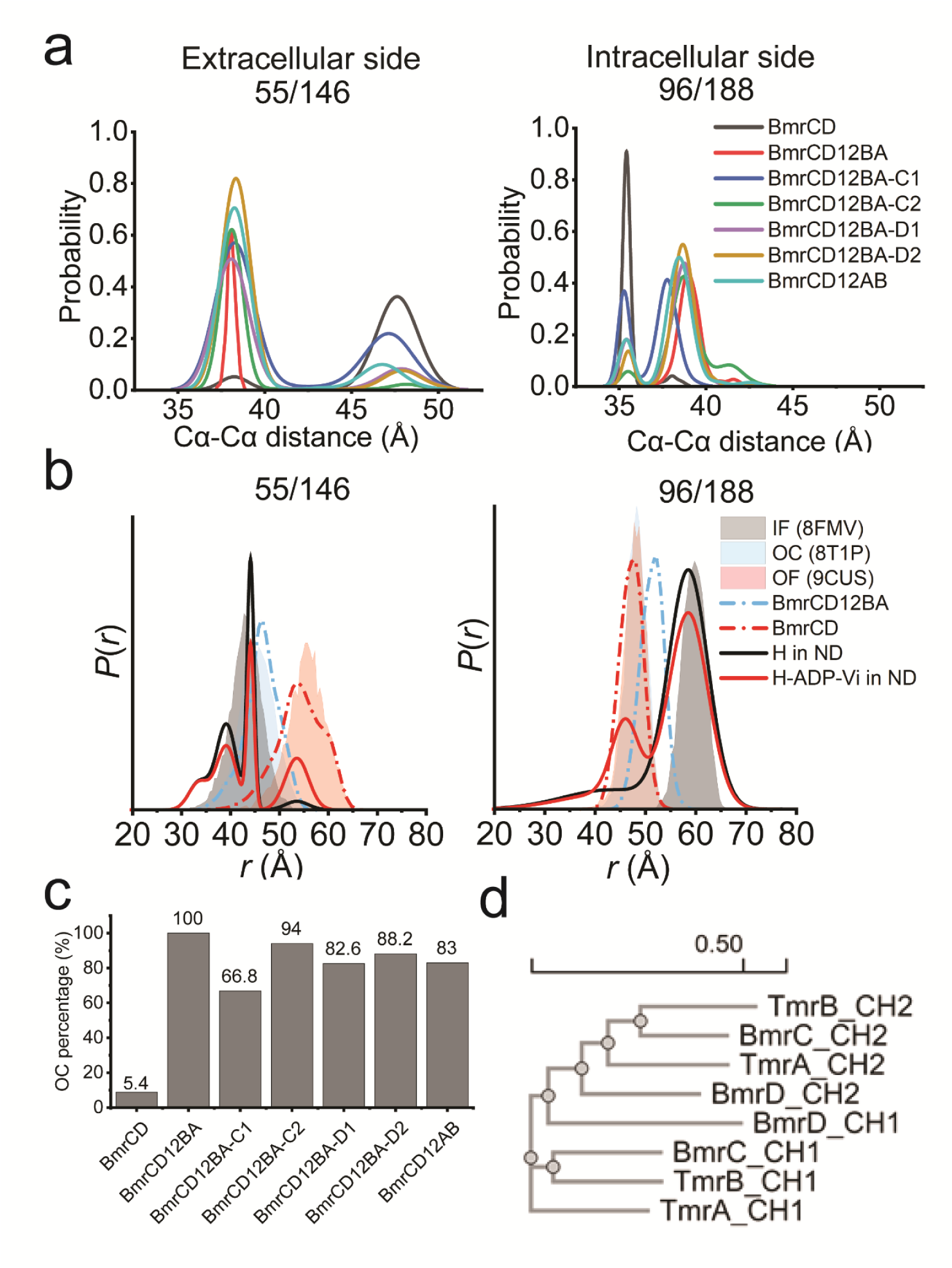
Replacement of the coupling helices in BmrCD increases the population of the OC conformation. **(a)** Cα-Cα distance measurements for the residue pairs on the extracellular (55^BmrC^/146^BmrD^) and intracellular (96^BmrC^/188^BmrD^) sides of BmrCD chimera with replaced coupling helices. **(b)** Validation of the chimeric BmrCD models. The experimental DEER distributions (solid lines) are compared to the simulations from ChiLife models (dashed lines) and from cryo-EM structures via MDDS (shaded areas). **(c)** Histogram of the percentage of OC conformation in the chimeric BmrCD. **(d)** Phylogenetic analysis of the coupling helices from BmrCD and TmrAB. Multiple alignment and phylogram construction were performed using Clustal W. The scale bar indicates the substitution rate per site.

In contrast, similar CHs replacement predictions conducted on TmrAB, where the CHs of TmrAB were substituted by the CHs of BmrCD, appeared to stabilize the OF conformations. Surprisingly, TmrAB12DC (with TmrA’s CHs of replaced by BmrD’s CH and TmrB’s by BmrC’s CH) as well as the opposite chimera TmrAB12DC showed a noticeable shift compared to WT with a detectable population of OF conformation, while TmrAB12CD (TmrA’s CHs replaced by BmrC’s and TmrB’s by BmrD’s) exhibited minimal changes (Fig. 10). The fact that the OF population is only a fraction of the model may suggest that there are sequence determinants of its stability in other segments of the transporter.

**Fig. 10.**
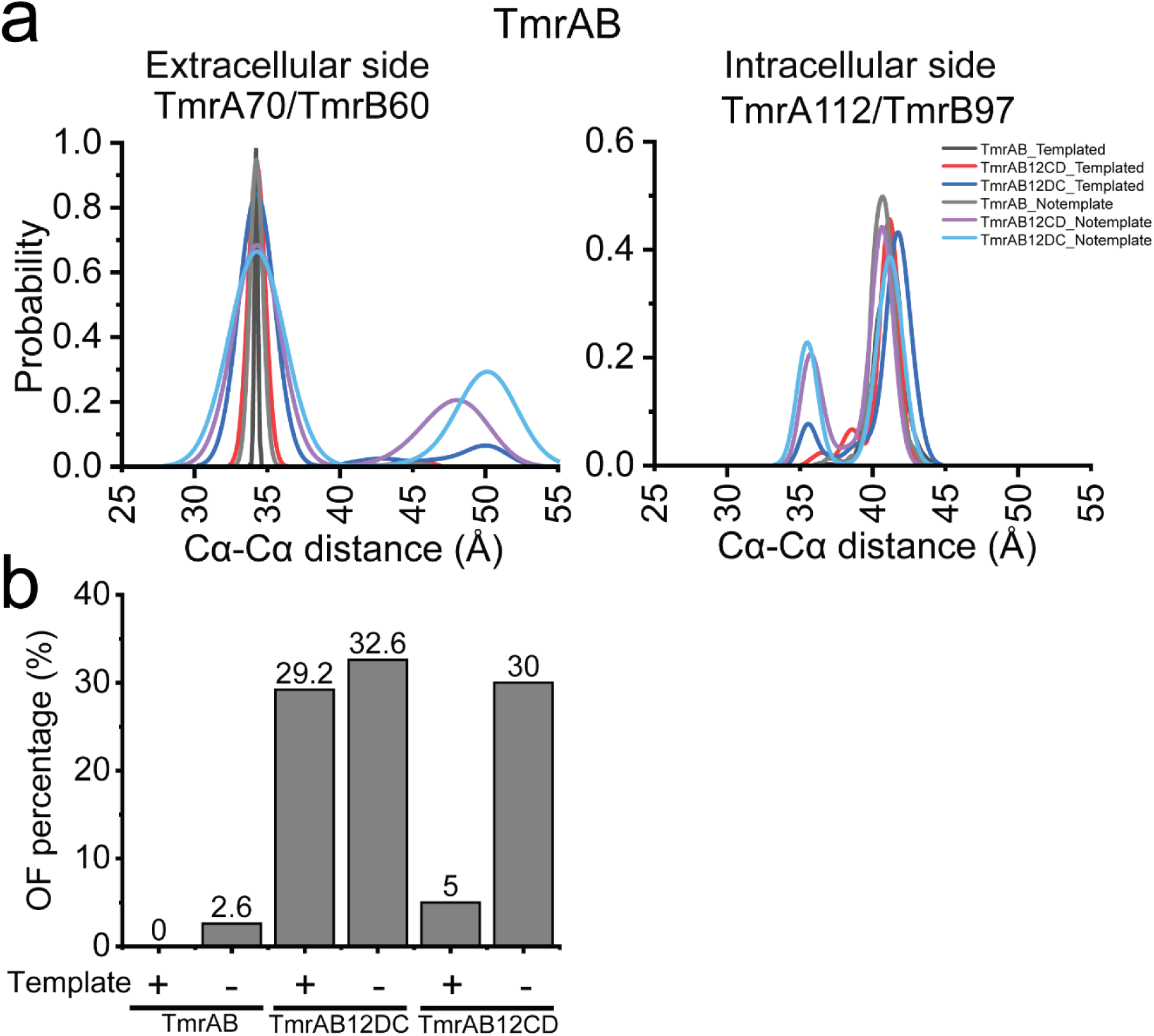
Replacement of the coupling helices in TmrAB sampled the OF state. **(a)** Cα-Cα distance measurements for the residue pairs on the extracellular (70^TmrA^/60^TmrB^) and intracellular (112^TmrA^/97^TmrB^) sides of TmrAB chimera with replaced coupling helices. **(b)** Histogram of the percentage of OF conformation in the chimeras of TmrAB.

### Concluding remarks

A number of interesting themes regarding AF3 predictions emerge from the focused analysis above. We find that AF3 can predict alternate protein conformations expected in the presence of ligands. The effects of ATP were not uniform across all four transporters suggesting that the predictions are context specific. In fact, we observed heterogenous predictions that seem congruent with experimental energetic findings. Furthermore, the predicted models did not simply reproduce experimental structures consistent with AF3 not being guided exclusively by templates. There were important correlations in the degree of implied disorder in the models with experimental findings. The extent to which these conclusions can be generalized remain to be investigated but suggest that AF3 provides a robust tool to test how ligand binding shifts protein conformational equilibria.

## MATERIALS AND METHODS

### Structure prediction by AF3

Structure prediction runs were executed using AF3 (*6*) (https://golgi.sandbox.google.com). For the four wild-type ABC transporters’ structure prediction, the input protein sequences for BmrCD are BmrC and BmrD, each with one copy; for TmrAB are TmrA and TmrB, each with one copy; for Pgp (human ortholog unless stated otherwise), with one copy; for MsbA, with two copies. The initial six screening conditions are apo (without any other component), 1ATP (with one copy of ATP as the ligand), 2ATP (with two copies of ATP as the ligands), 1Mg^2+^/2ATP (with two copies of ATP as the ligands, and one Mg^2+^ as the ion), 2Mg^2+^/2ATP (with two copies of ATP as the ligands, and two copies of Mg^2+^ as the ions), and 2Mg^2+^/1ATP1AD (with one copy of ATP and one copy of ADP as the ligands, and two copies of Mg^2+^ as the ions) in autoseed mode. For the engineered protein structure prediction, the wild-type protein sequence was replaced with the engineered one, as well as 2 copies of ATP and 2 copies of Mg^2+^ in autoseed mode. All jobs were repeated 100 times with random seeds, with templates applied unless otherwise noted.

### Predicted models clustered by Foldseek

To enable fast and sensitive clustering of the large number of models predicted by AF3, we applied Foldseek(*49*). For the monomeric human Pgp protein, we used monomer cluster function. Specifically, for its apo form, 500 models were clustered using default parameters with 30% minimum sequence identity, 50% minimum aligned residues, and generated a single cluster representing the inward-facing conformation. Similarly, a single cluster of the 2Mg^2+^/2ATP Pgp were obtained with default parameters, which showed the outward-facing conformation.

For BmrCD, TmrAB, and MsbA, the multimercluster function was applied with default parameters (multimer-tm-threshold: 0.65, chain-tm-threshold: 0.5, interface-lddt-threshold: 0.65) unless otherwise noted. The BmrCD IF, occluded, and OF conformations were selected from the apo, 1Mg^2+^/2ATP, 2Mg^2+^/2ATP conditions, respectively. MsbA IF and OF conformations were selected from the apo and 2ATP conditions, and TmrAB IF and OC conformations were selected from apo and 2Mg^2+^/2ATP conditions. Furthermore, The Cα-Cα distance distribution of intercellular pair 96/188 in 1Mg^2+^/2ATP BmrCD revealed five distinct populations: OF, semi OF, OC, asymmetric OC, and IF (Table 3). The representative models for OF, semi OF, OC, asymmetric OC, and IF were identified by Foldseek with default parameters.

MsbA occluded representative model was selected by multimercluster with a modified interface-lddt-threshold of 0.96 (and multimer-tm-threshold: 0.65, chain-tm-threshold: 0.5). Three clusters were generated from 500 models, and 9 of them are occluded. Since AF3 could not predict an OF conformation for wildtype TmrAB, we used a chimeric TmrAB with its coupling helices replaced by those from BmrCD (TmrAB12DC), two clusters were produced by multimercluster with strict parameters (0.95 for multimer-tm-threshold, chain-tm-threshold, and interface-lddt-threshold) from 500 models, 52 of which exhibited the OF conformation.

All the presentative models are shown in cartoon representation in Fig. 2. The TM-scores and RMSD values between cryo-EM structures and the representative AF3 models are listed in Table 1.

### Root Mean Square Deviation (RMSD) and Root Mean Square Fluctuation (RMSF) analysis

**RMSD** measures the average distance between corresponding atoms in two superimposed protein structures. It is defined as:

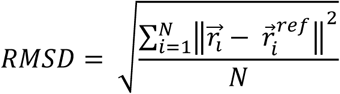

Where: RMSD is the Root Mean Square Deviation of *N* atoms; 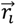 and 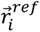 are the coordinate vectors of the *i*-th atom in the target and reference structures after superposition; *N* is the number of atoms used for alignment. 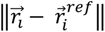 is the Euclidean distance between the two corresponding atoms.

**RMSF** quantifies how much each residue’s position fluctuates around its average position in a protein structure ensemble. For RMSF analysis, all predicted models were converted from CIF to PDB format. The first model was used as the reference structure, and Cα atoms were extracted for alignment based on overall structure. Subsequently, the average position of each atoms was calculated and used as the reference point for computing the root mean square fluctuation across the entire set of models. It is defined as:

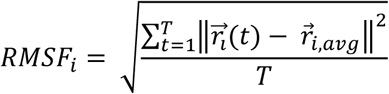

Where: *RMSF_i_* is the Root Mean Square Fluctuation of atom *i*; 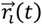 is the coordinate vector of atom *i* in model *t*; 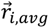 is the average coordinate vector of atom *i*; *T* is the total number of models under one condition). 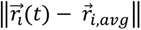 is the Euclidean distance between atom *i* in model *t* and its average position.

### ChiLife used for *in silico* spin labeling

Given the hundreds of models generated by AF3 in our study, we investigated the simulated distance distributions of the IF, OC, and OF conformation ensembles. The results were compared to distance distributions simulated by MDDS for cryo-EM structures, as well as experimental DEER data.

Briefly, the chiLife Python package(*50*) was applied to calculate ensemble distance distribution through *in silico* modeling of site-directed spin labels (SDSL). The class method xl.SpinLabel.from_mmm was employed to create spin label objects using data generated by the MATLAB program MMM (Multiscale Modeling of Macromolecules), which applies a rotamer library approach to predict spin label conformations. We utilized the common nitroxide spin label R1M along with MDAnalysis universe objects for site-directed spin labeling and molecular dynamics trajectories analysis.

The extracellular spin labeling sites for BmrCD, MsbA, and Pgp are 55^BmrC^/146^BmrD^, 158/158, 96/749, respectively; while the intracellular sites were 96 ^BmrC^/188 ^BmrD^, 103/103, 149/791, respectively. The conditions used for chiLife analysis of BmrCD include apo, 1Mg^2+^/2ATP, and 2Mg^2+^/2ATP; for Pgp, the conditions were apo and 2Mg^2+^/2ATP; for MsbA, the conditions were apo, 1ATP, and 2ATP. For each condition, 500 models were used for simulated distance distribution calculations, which were normalized by probability and plotted in Fig. 5 (shown as dashed lines).

### Coupling helices switch of BmrCD and TmrAB

Through sequence and structure analysis of BmrCD and TmrAB, the sequences of coupling helices for BmrC are defined as TMSPPFYEKNRT (residue 103-114, denoted as C1) and SGVRVIRAYVQ (residue 206-216, denoted as C2); the sequences of coupling helices for BmrD are KMPIRYFDNLPA (residue 195-206, denoted as D1) and QGMTIIQAFRH (residue 297-307, denoted as D2); the sequences of coupling helices for TmrA are RLHPGFYDRNPV (residue 119-130, denoted as A1) and SGVETIQLFVK (residue 221-231, denoted as A2); the sequences of coupling helices for TmrB are TLDRDFYHKHRV (residue 104-115, denoted as B1) and SGIRVVKGYAL (residue 206-216, denoted as B2), respectively. BmrCD12BA refers to BmrCD where the coupling helices C1 and C2 were replaced with B1 and B2, respectively; and the coupling helices D1 and D2 were replaced with A1 and A2, respectively. Meanwhile, BmrCD12BA-C1 is BmrCD12BA, but with C1 unchanged. Similarly, BmrCD12BA-C2, BmrCD12BA-D1, BmrCD12BA-D2 were engineered based on BmrCD12BA, but with C1, D1, and D2 unchanged respectively. BmrCD12AB refers to BmrCD with the coupling helices C1 and C2 replaced by A1 and A2, respectively while D1 and D2 were replaced by B1 and B2, respectively (see Table 4 for more details).

## Author contributions

Q.T. prepared all input JSON files for AF3 predictions, performed all AF3 predictions for chimeric proteins, conducted non-templated predictions using the online AF3 server, carried out all data analysis, and prepared the figures. T.W. performed the subsampling. B.S. conducted template-based AF3 predictions for the wild-type type IV ABC transporter using a local installation. Q.T. and H.S.M. designed the research, analyzed and interpreted the results, and wrote the manuscript with input from all authors.

## Funding

This work was supported by National Institutes of Health grant 3R35GM152382 to H.S.M.

## Competing interests

Authors declare that they have no competing interests.

## Notes

### Competing Interest Statement

The authors have declared no competing interest.

