## Supplemental information for "Prediction of ligand-dependent conformational sampling of ABC transporters by AlphaFold3 and correlation to experimental structures and energetics"

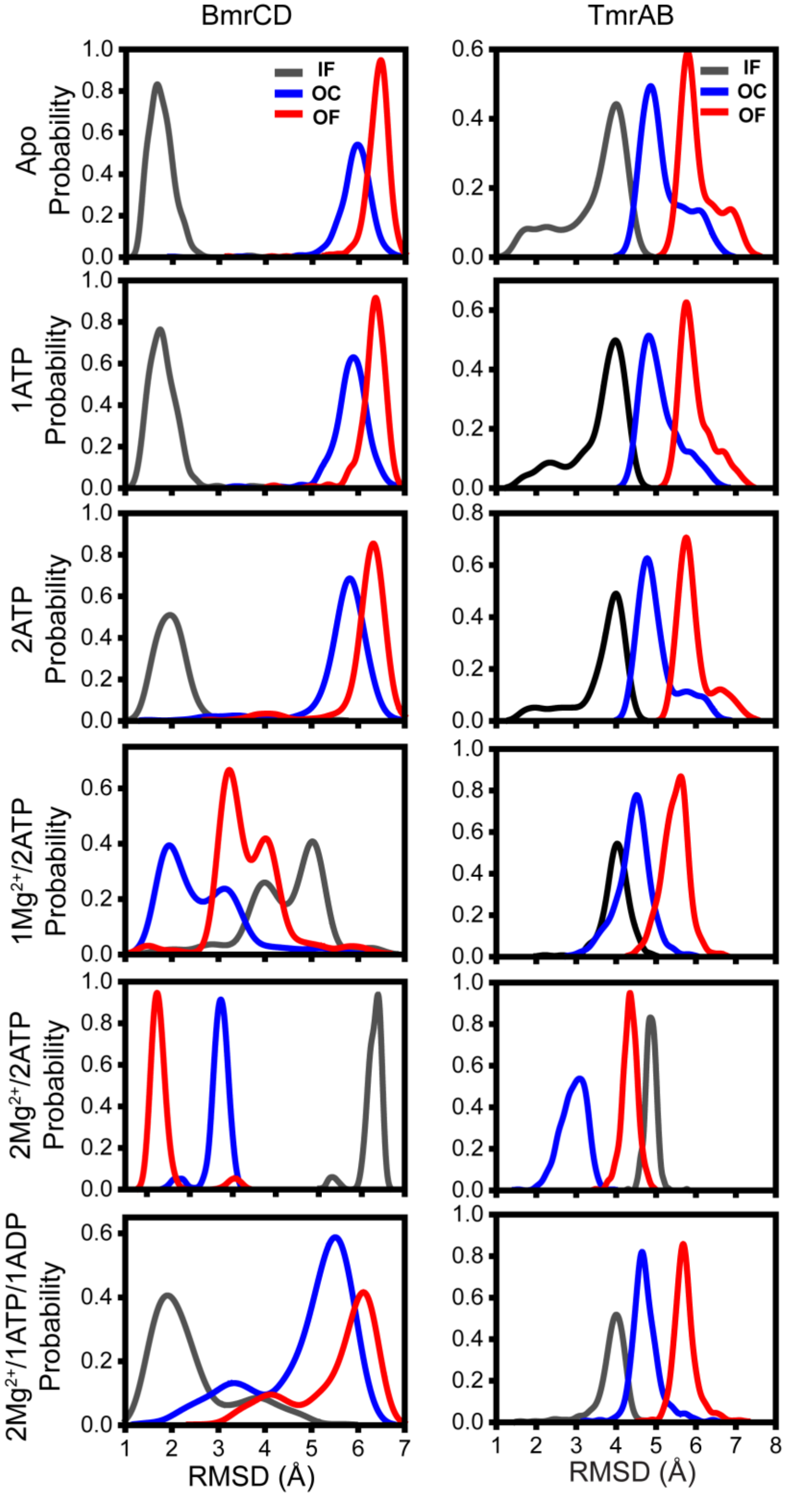


**Fig. S1 RMSD analysis of AF3 predictions for BmrCD and TmrAB in the presence of combinations of nucleotide and $Mg^{2+}$.**

AF3 predictions were performed for BmrCD and TmrAB under six nucleotide conditions: apo, 1ATP, 2ATP, 1$Mg^{2+}$/2ATP, 2$Mg^{2+}$/2ATP, and 2$Mg^{2+}$/1ATP1ADP. For each condition, 500 models were generated using random seeds. RMSD values for the models in each condition were calculated with MMalign by aligning them to the reference cryo-EM structures of IF, OC, and OF states. The resulting RMSD distributions for the IF, OC, and OF are shown in grey, blue and red, respectively. Reference PDB IDs: BmrCD (IF: 8FMV, OC: 8T1P, OF: 9CUS), TmrAB (IF: 6RAN, OC: 6RAI, OF: 6RAH).

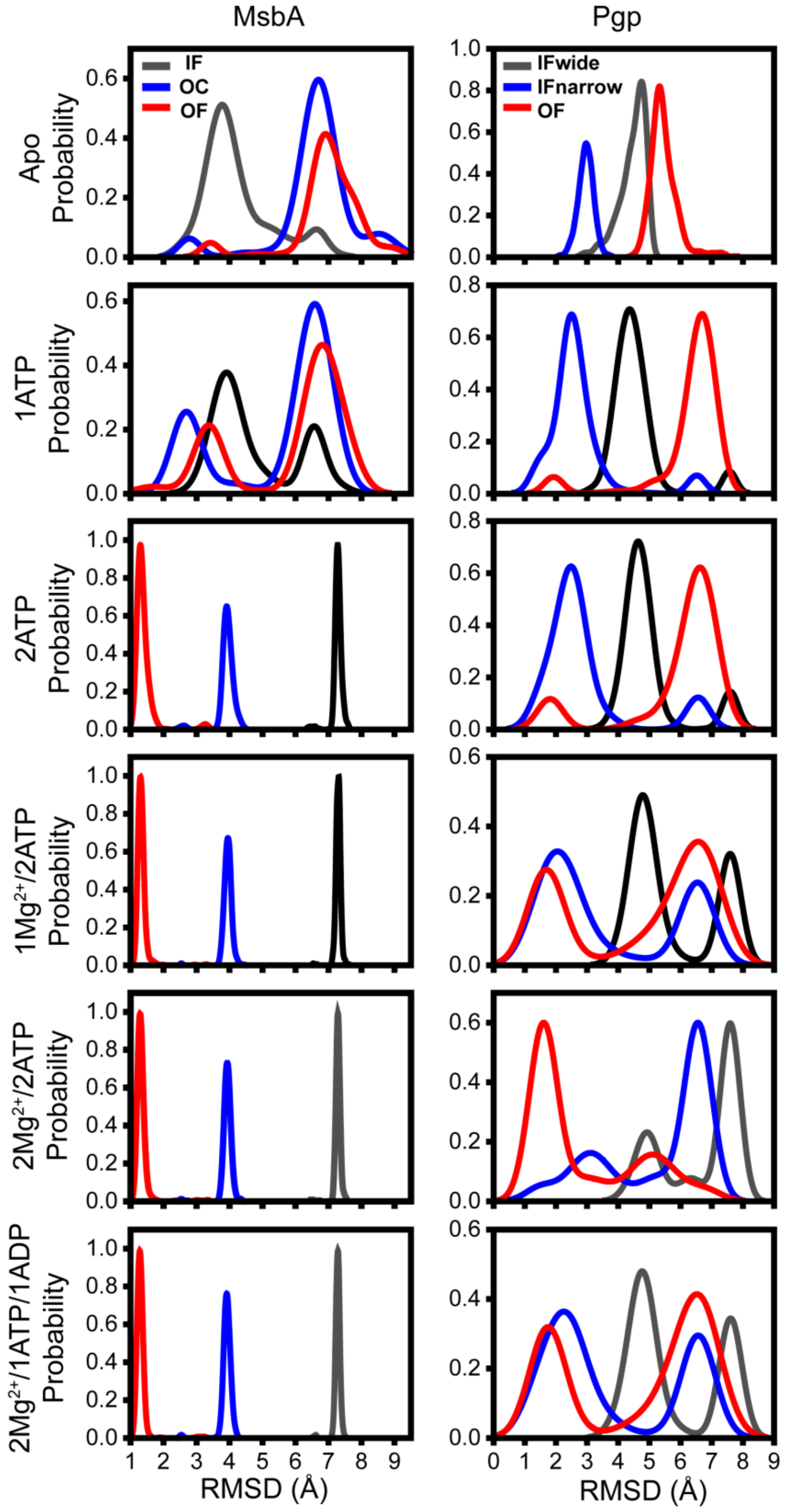


**Fig. S2 RMSD analysis of AF3 predictions for MsbA and Pgp in the presence of combinations of nucleotide and $Mg^{2+}$.**

AF3 predictions were performed for MsbA and Pgp under apo, 1ATP, 2ATP, 1$Mg^{2+}$/2ATP, 2$Mg^{2+}$/2ATP, and 2$Mg^{2+}$/1ATP1ADP. For each condition, 500 models were generated using random seeds. RMSD values for the models in each condition were calculated by aligning them to the reference cryo-EM structures of IF, OC, and OF states using MMalign for MsbA and TMalign for Pgp. The resulting RMSD distributions for the IF, OC, and OF are shown in grey, blue and red, respectively. Reference PDB IDs: MsbA (IF: 5TV4, OC: 5TTP, OF: 8DMM), Pgp (IF-wide: 3G5U, IF-narrow: 6QEX, OF: 6C0V).

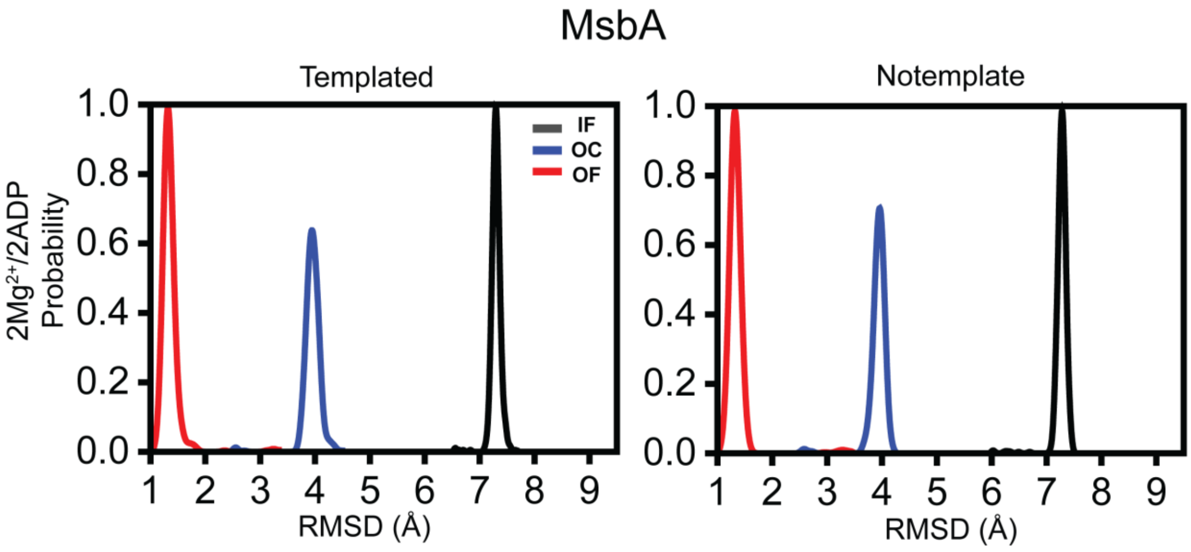


**Fig. S3 RMSD analysis of AF3 models with and without templates for MsbA under $2Mg^{2+}$/2ADP.**

AF3 was used to predict MsbA structures under $2Mg^{2+}$/2ADP with (Templated) and without (Notemplate) template. For each condition, 500 models were generated using random seeds. RMSD values for the models in each condition were calculated by aligning them to the reference cryo-EM structures of IF (5TV4), OC (5TTP), and OF (8DMM) states using MMalign for MsbA. The resulting RMSD distributions for the IF, OC, and OF are shown in grey, blue and red, respectively.

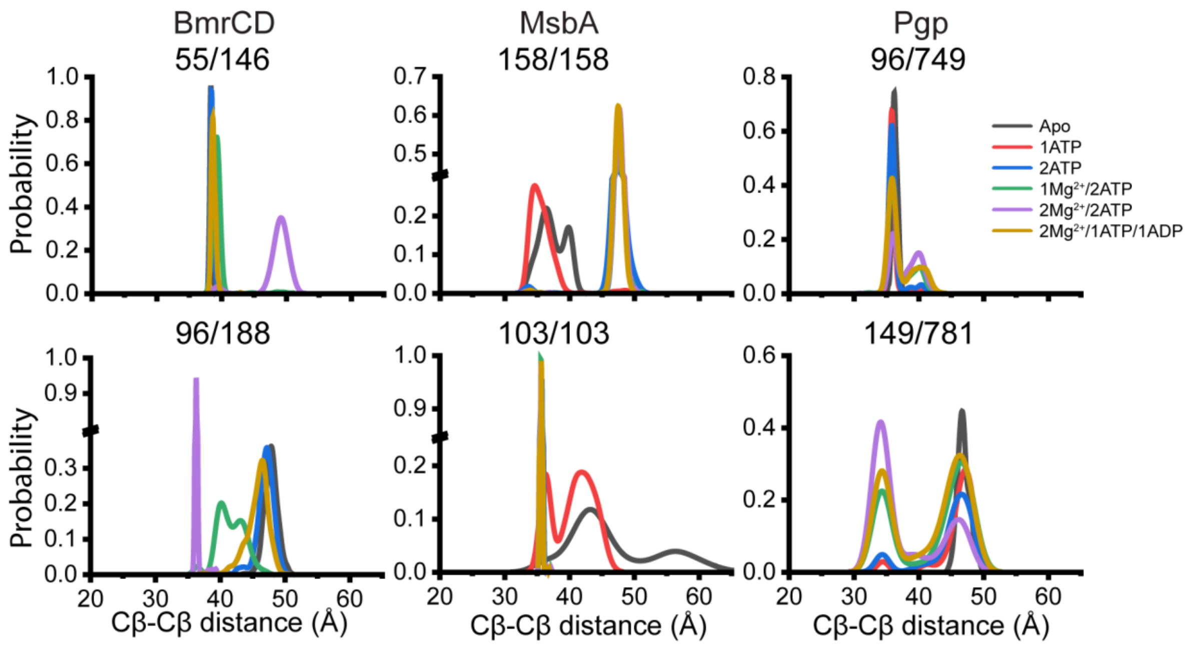


**Fig. S4 Distinct conformational transitions revealed by the Cβ-Cβ distance analysis for BmrCD, MsbA, and Pgp.**

Distribution plots of the Cβ-Cβ distances of the residue pairs on the extracellular side (upper panels) and intracellular side (lower panels) of BmrCD (left panels), MsbA (middle panels), and Pgp (right panels). Data for six ligand conditions (apo, 1ATP, 2ATP, 1$Mg^{2+}$/2ATP, 2$Mg^{2+}$/2ATP, and 2$Mg^{2+}$/1ATP1ADP) are plotted for each pair. Distribution analysis was performed with Origin software (bin size equals to 1 Å; data height is calculated by Relative Frequency).

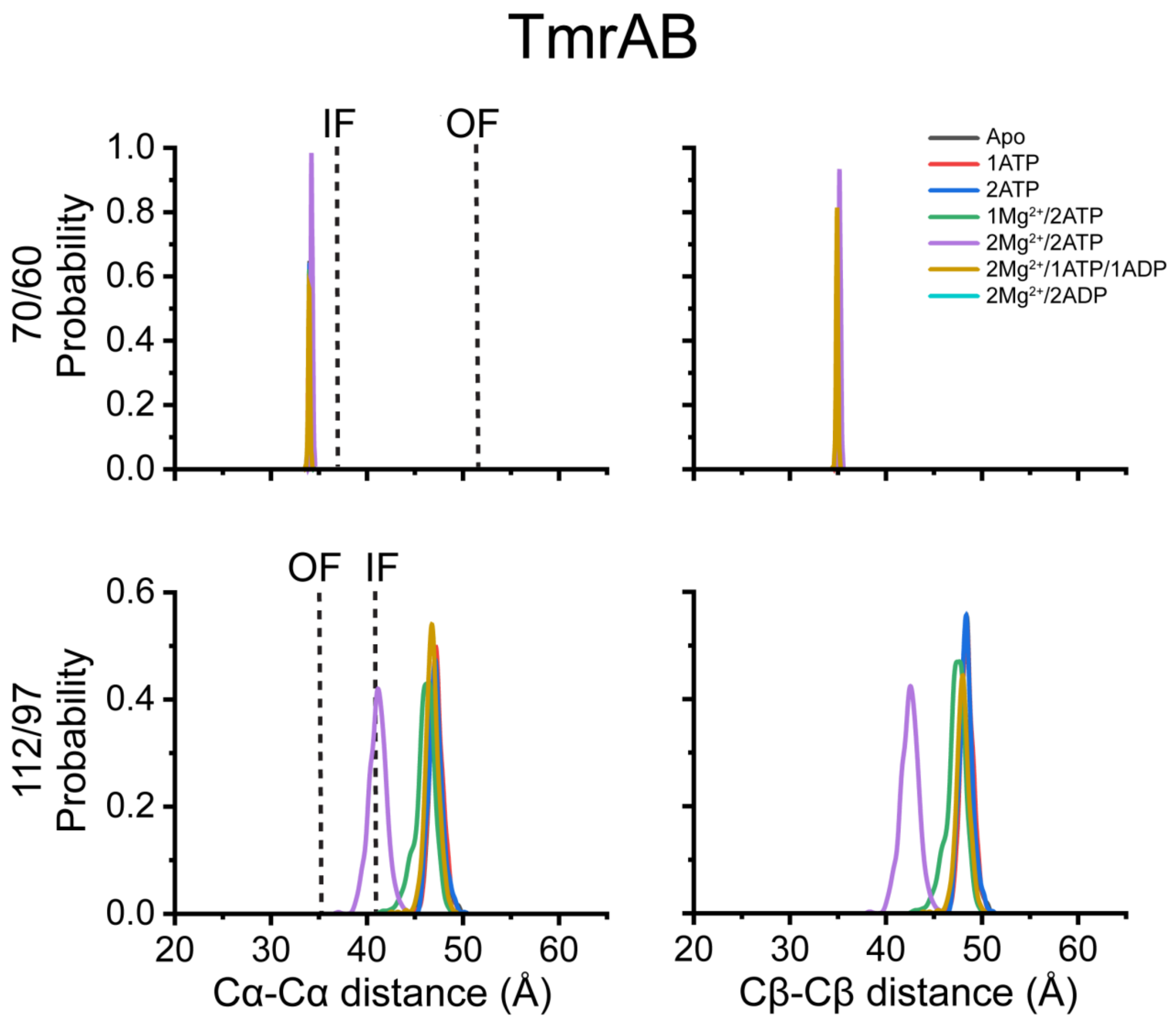


**Fig. S5 Distinct conformational transitions in TmrAB revealed by the Cα-Cα and Cβ-Cβ distances analysis.**

Distribution plots of the Cα-Cα (left panels) and Cβ-Cβ (right panels) distances of the residue pairs on the extracellular side (upper panels) and intracellular side (lower panels) of TmrAB. Residue pair 70/60 and 112/97 are denoted for $70^{TmrA}/60^{TmrB}$ and $112^{TmrA}/97^{TmrB}$. Data for ligand conditions (apo, 1ATP, 2ATP, $1Mg^{2+}/2ATP$, $2Mg^{2+}/2ATP$, $2Mg^{2+}/1ATP1ADP$ and $2Mg^{2+}/2ADP$) are plotted for each pair. The dashed lines indicate the distance derived from the cryo-EM structures of the corresponding protein in its IF and OF conformations. Distribution analysis was performed with Origin software (bin size equals to 1 Å; data height is calculated by Relative Frequency).

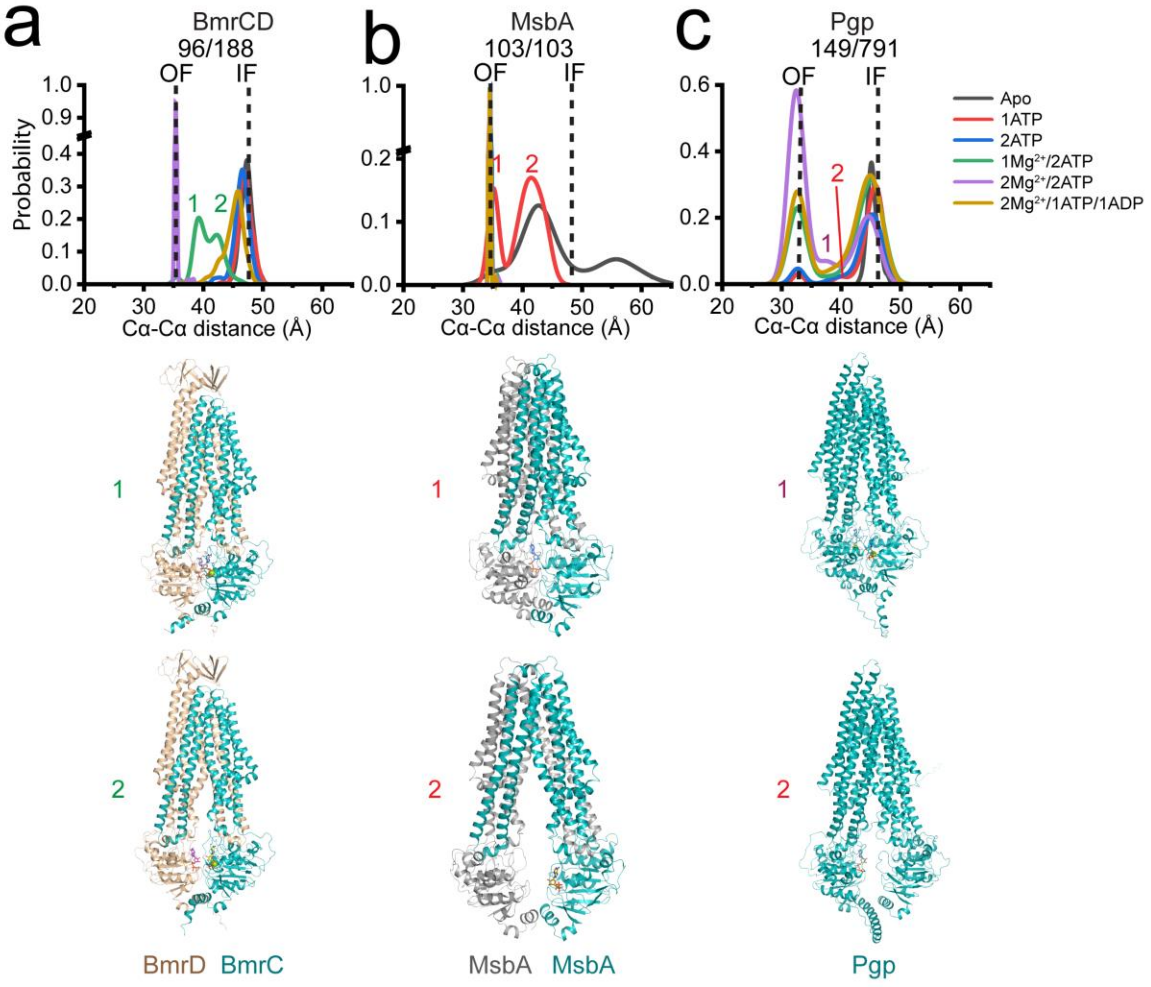


**Fig. S6 Identifying intermediate states through intracellular Cα-Cα distance distributions in BmrCD, MsbA, and Pgp.**

Distribution of intracellular Cα-Cα distances of the residue pairs in BmrCD (left panels), MsbA (middle panels), and Pgp (right panels), which are derived from Fig. 6b. The respective peaks correspond to intermediate states are highlighted, and the representative structures are selected via Foldseek(*1*) and displayed as cartoon below each plot. The color scheme follows that of Fig. 2.

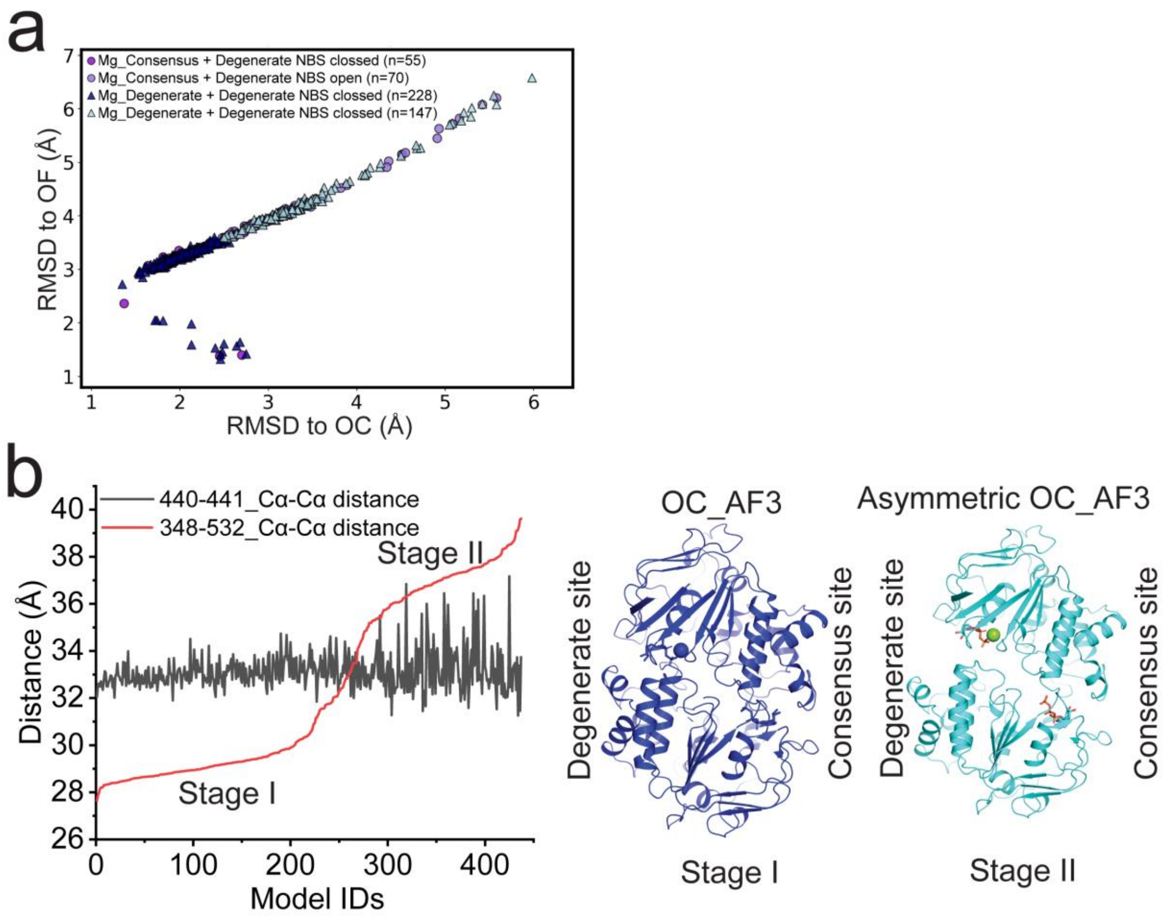


**Fig. S7 Asymmetric NBS states revealed by predicted BmrCD structures under $1Mg^{2+}$/2ATP. (a)** RMSD plot for the $1Mg^{2+}$/2ATP condition of BmrCD, highlighting the $Mg^{2+}$-binding site and degenerate NBS conformation. Models are marked with a circle ($Mg^{2+}$ bound at the consensus site) or a triangle ($Mg^{2+}$ bound at the degenerate site), and the fill color indicates the states of the degenerate NBS: dark (closed) or light (open). **(b)** Cα-Cα distance plot for the residue pairs in BmrCD nucleotide-binding sites (NBSs). The black line represents the consensus site pair $440^{BmrC}/441^{BmrD}$ (440-441Cα-Cα distances), while the red line represents the degenerate site pair $348^{BmrC}/532^{BmrD}$ (348-532 Cα-Cα distances). Two distinct stages can be identified for the degenerate site, and the representative model were selected via Foldseek(*1*) and are shown as cartoons, with $Mg^{2+}$ as spheres and ATP as sticks.

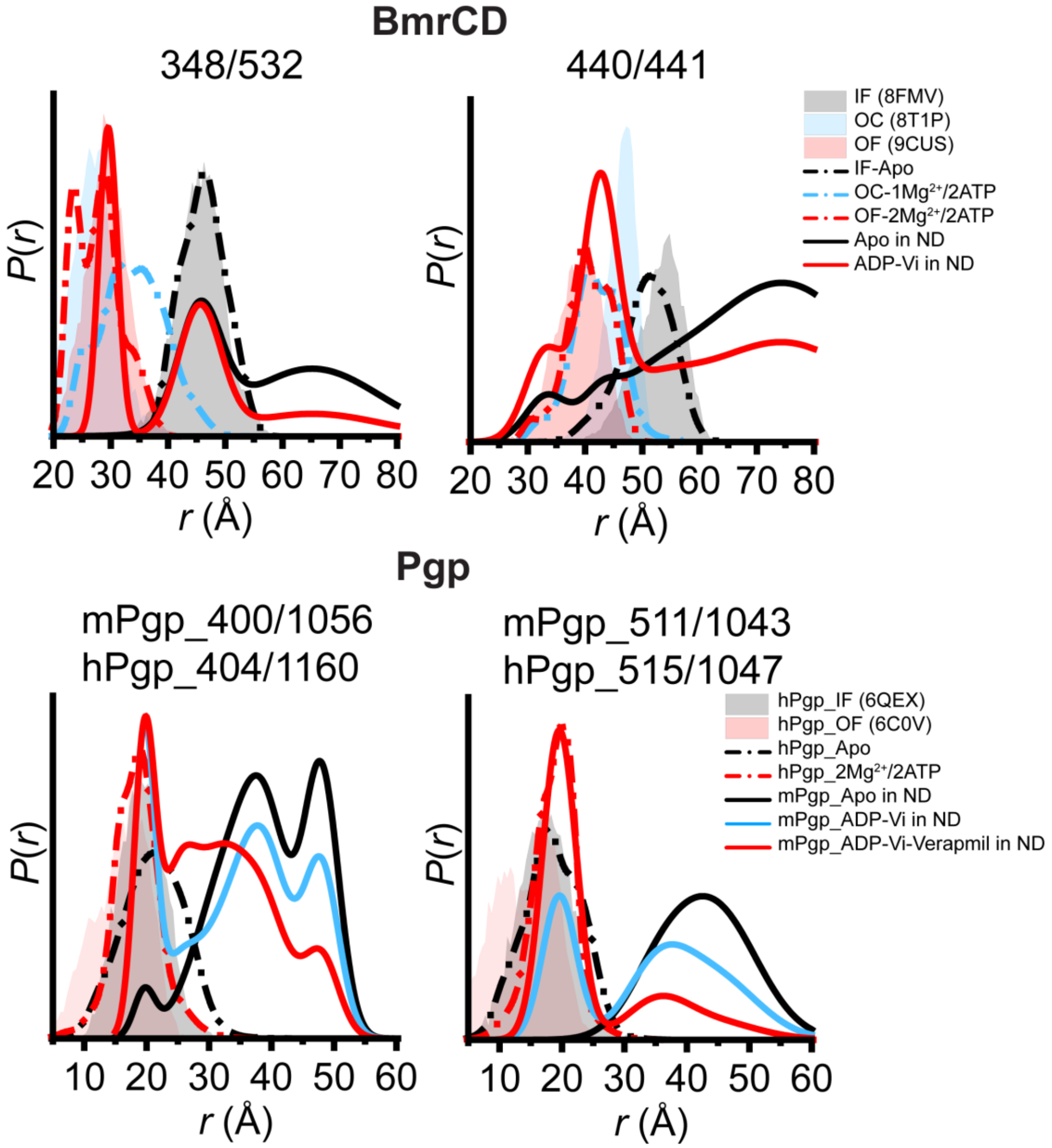


**Fig. S8 Asymmetric characteristics of BmrCd and Pgp NBSs revealed by AF3, cryo-EM structures and DEER spectroscopy.** AF3 models were validated by comparing simulated distance distributions to experimental DEER data for the spin-labeling sites at the NBS of BmrCD and Pgp. Spin-labeling pair 348/532 and 440/441 are denoted for $348^{BmrC}/532^{BmrD}$ and $440^{BmrC}/441^{BmrD}$, respectively. For each pair, the shaded area represents the distribution simulated from cryo-EM structure using MDDS. The dashed line shows the distribution simulated from the corresponding ChiLife(*2*) models, and the solid line shows the experimental DEER distribution.

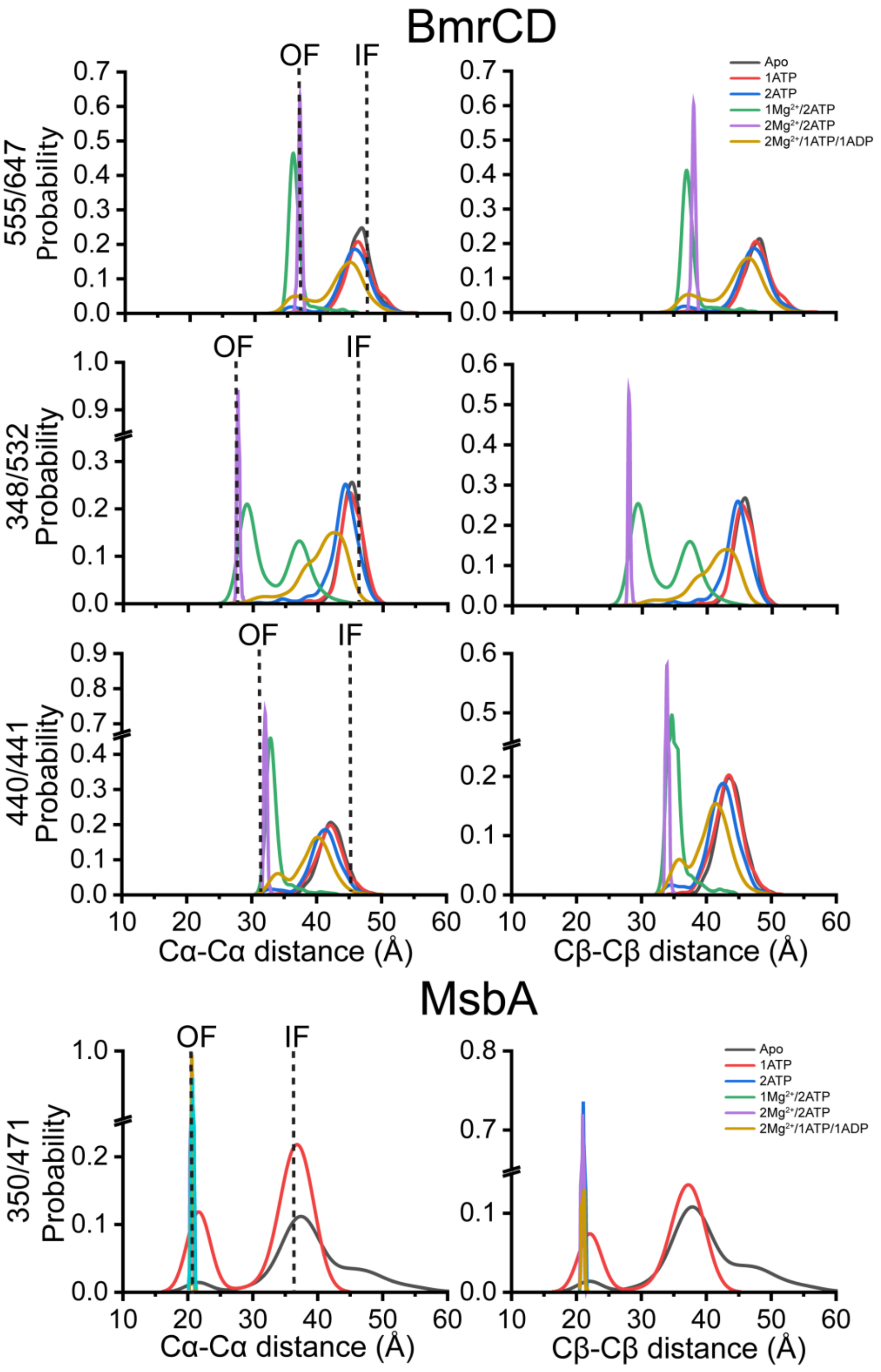


**Fig. S9 Cα-Cα and Cβ-Cβ distances analysis for the NBSs of BmrCD and MsbA.**

Distribution plots of the Cα-Cα (left panels) and Cβ-Cβ (right panels) distances for the residue pairs at the NBSs of BmrCD and MsbA. Residue pair 348/532, 440/441 and 555/647 are denoted for $348^{BmrC}/532^{BmrD}$, $440^{BmrC}/441^{BmrD}$ and $555^{BmrC}/647^{BmrD}$. Data for ligand conditions (apo, 1ATP, 2ATP, $1Mg^{2+}/2ATP$, $2Mg^{2+}/2ATP$, and $2Mg^{2+}/1ATP1ADP$) are plotted for each pair. The dashed lines indicate the distance derived from the cryo-EM structures of the corresponding protein in its IF and OF conformations. Distribution analysis was performed with Origin software (bin size equals to 1 Å; data height is calculated by Relative Frequency).

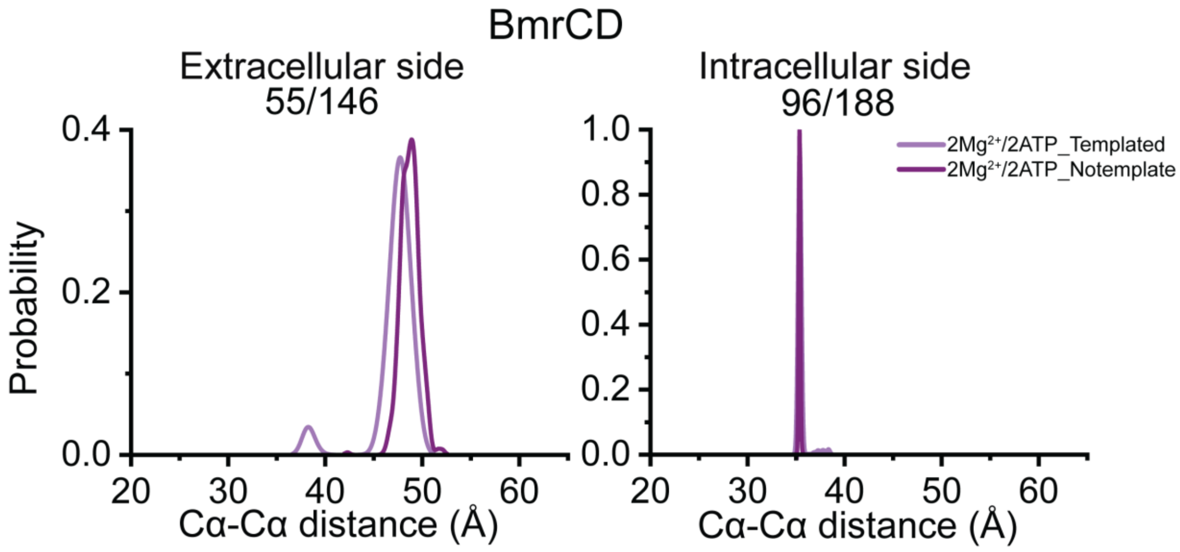


**Fig. S10 Comparison of AF3 predictions for BmrCD with and without templates.**

Cα-Cα distance measurements for the residue pairs on the extracellular 55/146 ($55^{BmrC}/146^{BmrD}$) and intracellular 96/188 ($96^{BmrC}/188^{BmrD}$) sides of BmrCD with template (Templated) or without template (Notemplate) in the presence of $2Mg^{2+}/2ATP$.
